# Architecture of the yeast Pol III pre-termination complex and pausing mechanism on poly-dT termination signals

**DOI:** 10.1101/2022.02.28.482286

**Authors:** Mathias Girbig, Juanjuan Xie, Helga Grötsch, Domenico Libri, Odil Porrua, Christoph W. Müller

## Abstract

RNA polymerase (Pol) III is specialized to transcribe short, abundant RNAs, for which it terminates transcription on poly-thymine (dT) stretches on the non-template (NT) strand. When Pol III reaches the termination signal, it pauses and forms the pre-termination complex (PTC). Here, we report cryo-EM structures of the yeast Pol III PTC and complementary functional states at 2.7-3.9 Å resolution. Pol III recognizes the poly-dT termination signal with subunit C128 that forms a hydrogen-bond network with the NT-strand and, thereby, induces pausing. Mutating key interacting residues interferes with transcription termination *in vitro*, impairs yeast growth, and causes global termination defects *in vivo* confirming our structural results. Additional cryo-EM analysis reveals that C53-C37, a Pol III subcomplex and key termination factor, participates indirectly in Pol III termination. We propose a mechanistic model of Pol III transcription termination and rationalize why Pol III, unlike Pol I and Pol II, terminates on poly-dT signals.

## INTRODUCTION

Eukaryotic RNA transcription is carried out by up to five DNA-dependent RNA polymerases (Pols), of which Pol I, Pol II, and Pol III are central to all eukaryotes. Pol I transcribes ribosomal RNA (rRNA). Pol II produces messenger RNAs (mRNAs) and various non-coding RNAs. Pol III synthesizes very short – but highly abundant RNA molecules – such as transfer RNAs (tRNAs), the U6 small nuclear RNA (snRNA), and the 5S rRNA. Pol III-transcribed RNAs contribute the vast majority of mature RNA molecules to the pool of cellular RNA with tRNAs being the most abundant transcripts (Palazzo and Lee, 2015). Pol III is highly specialized to cope with the high demand of its target transcripts and, as such, requires much less additional factors to initiate, elongate, and terminate transcription to streamline these processes.

With 17 subunits, Pol III is the largest eukaryotic RNA polymerase. Pol III contains a conserved 10-subunit catalytic core and a 2-subunit stalk domain, which are both also present in Pol I and Pol II. In addition, it features the C53-C37 heterodimer and the C82-C34-C31 heterotrimer. The C53-C37 heterodimer shares structural homology with the Pol II general transcription factor (GTF) TFIIF (Geiger et al., 2010; Hoffmann et al., 2015). The C82-C34-C31 heterotrimer subunits are partly homologous to TFIIE and TFIIF (Abascal-Palacios et al., 2018; Lefèvre et al., 2011; Vannini and Cramer, 2012; Vorländer et al., 2018). Pol III also harbors the RNA-cleaving subunit C11 (RPC10 in humans) whose C-terminal domain is homologous to the RNA-cleaving domain of the Pol II elongation factor TFIIS (Chedin et al., 1998). Similarly to TFIIF and TFIIE, C53-C37 and C82-C34-C31 are both required to initiate transcription (Brun, 1997; Kassavetis et al., 2010; Landrieux et al., 2006; Lefèvre et al., 2011; Thuillier et al., 1995; Wang and Roeder, 1997). In addition, the C53-C37 heterodimer and C11 are both crucial for Pol III transcription termination (Arimbasseri and Maraia, 2013, 2015; Chedin et al., 1998; Landrieux et al., 2006; Mishra and Maraia, 2019; Mishra et al., 2021; Rijal and Maraia, 2013).

Transcription termination is the final step in the transcription cycle and involves the release of both the transcribed RNA and the DNA so that the RNA polymerase gets freed-up and is, potentially, available for a new round of transcription. In the Pol I and Pol II systems, a multitude of accessory factors are needed for transcription termination (Porrua and Libri, 2015; Richard and Manley, 2009). On the contrary, Pol III relies on fewer elements to terminate transcription, which is a key specialization of the Pol III system. The precise termination process allows fast recycling of Pol III to a new round of transcription in a process termed facilitated recycling (Dieci and Sentenac, 1996). Pol III terminates transcription on sites that contain a continuous stretch of thymine bases (dT) on the non-template (NT) strand of the DNA (Allison and Hall, 1985; Bogenhagen and Brown, 1981; Cozzarelli et al., 1983; Watson et al., 1984). Yeast Pol III requires at least 5 dTs to correctly terminate transcription (Allison and Hall, 1985) and, *in vitro*, does not rely on any external factors to do so (Braglia et al., 2005; Landrieux et al., 2006). The Pol III termination mechanism involves an intricate network of interactions of multiple Pol III subunits, the NT-strand, and the DNA:RNA hybrid in the Pol III active site that work hand-in-hand to enable precise and efficient termination. Pol III utilizes its in-built subunits C160 (Rijal and Maraia, 2016), C128 (James and Hall, 1990; Shaaban et al., 1995, 1996) and C53-C37 (Arimbasseri and Maraia, 2013; Landrieux et al., 2006; Rijal and Maraia, 2013) to facilitate termination on poly-dT signals. Subunit C11 is also necessary for Pol III transcription termination – which does not require its RNA-cleaving activity (Chedin et al., 1998; Iben et al., 2011; Landrieux et al., 2006) and is critical for the facilitated recycling process (Landrieux et al., 2006; Mishra et al., 2021).

Prior to termination, Pol III has to reduce its transcription rate (Rijal and Maraia, 2016), to which the C53-C37 heterodimer contributes (Arimbasseri and Maraia, 2013). When Pol III encounters the poly-dT termination sequence, it pauses on the 4^th^ dT, thus forming the pre-termination complex (PTC) (Arimbasseri and Maraia, 2015). It was proposed that Pol III undergoes conformational changes upon PTC formation, which mediate Pol III pausing and prime it for subsequent termination once it encounters the 5^th^ dT (Arimbasseri and Maraia, 2015). The C53-C37 heterodimer was shown to be critical for PTC formation (Arimbasseri and Maraia, 2015). In the cryogenic electron microscopy (cryo-EM) structure of the Pol III elongation complex (EC), C53-C37 binds the Pol III core near the region where the unwound NT-strand presumably is located (Hoffmann et al., 2015). This observation led to the hypothesis that a flexible loop in subunit C37 (aa 197-224) recognizes the termination signal and, thereby, drives PTC formation (Hoffmann et al., 2015). Since the unwound NT-strand in the Pol III EC structure was, however, only poorly resolved (which is also the case for most Pol I and Pol II structures), the exact path of the NT-strand remained elusive. It is, therefore, unclear how Pol III recognizes the poly-dT termination signal in the context of the Pol III PTC and how this facilitates pausing of Pol III and predisposes it for the termination process.

Pol III has been long thought to terminate in the absence of any cofactors. Recently though, the helicase Sen1 was shown to interact with Pol III and to also function in Pol III transcription termination (Rivosecchi et al., 2019; Xie et al., 2021). Deleting Sen1 in *S. pombe* cells resulted in increased termination RT products, which led the authors to conclude that Sen1 acts as a primary termination factor (Rivosecchi et al., 2019). A more recent study, in which a Sen1 mutant that failed to interact with Pol III in *S. cerevisiae* was investigated, also revealed that disrupting the Sen1-Pol III interaction causes increased termination RT of Pol III (Xie et al., 2021). However, the later study concluded that *S. cerevisiae* Sen1 more likely functions as a “fail-safe” termination factor that acts on secondary poly-dT termination sites residing downstream of the first termination signal (Xie et al., 2021). In that scenario, Sen1 would trigger the release of Pol III that failed to terminate on its primary termination site and got stuck on downstream poly-dT signals.

In the recent years, the eukaryotic transcription machineries have been extensively studied via cryo-EM, which greatly advanced our understanding of the mechanism of eukaryotic transcription. However, most of the structural investigations focused on how transcription initiation and elongation are facilitated and regulated. Consequently, transcription termination is the least understood stage in the eukaryotic transcription cycle. In the case of Pol III, structural insights could be obtained of initiating (Abascal-Palacios et al., 2018; Han et al., 2018; Vorländer et al., 2018) and elongating yeast Pol III (Hoffmann et al., 2015). In addition, we and others have recently reported the structures of human Pol III (Girbig et al., 2021; Li et al., 2021; Ramsay et al., 2020; Wang et al., 2021). However, structural information into the Pol III termination mechanism was largely missing, which strongly limited our understanding of the complete Pol III transcription cycle. A recent cryo-EM structure of the human Pol III PTC at 3.6 Å resolution showed that subunit RPC2 (the human ortholog to C128) recognizes the poly-dT termination sequence (Hou et al., 2021). However, to what extent the contact sites between RPC2 and the termination sequence are critical to termination is still not known due to a lack of functional validation of this interaction. Furthermore, the human Pol III PTC structure could not explain why the C53-C37 (RPC4-RPC5 in humans) is critical for correct termination.

Here, we report the cryo-EM structure of the yeast Pol III PTC at a resolution of 2.8 Å. The structure confirms that the yeast Pol III subunit C128 recognizes the poly-dT termination signal and explains how pausing is achieved, which we extensively validate with structure-function studies. Moreover, we present complementary high-resolution structures that shed light on how C53-C37 participates in transcription termination and how PTC formation may drive the subsequent termination process.

## RESULTS

### Purified yeast Pol III terminates transcription in vitro on poly-dT termination signals

To study the molecular mechanism of Pol III transcription termination on poly-dT sequences, we used the 17-subunit yeast Pol III complex. We first tested if the yeast Pol III sample, endogenously purified from *S. cerevisiae* fermenter cultures and used for earlier structural studies (Hoffmann et al., 2015; Vorländer et al., 2018, 2020), is capable to terminate transcription on poly-dT sequences as described earlier. We used a double-stranded (ds) DNA construct composed of the *S. cerevisiae* U6 snDNA gene promoter and followed by the 3’-flanking sequence of the *S. cerevisiae* tRNA gene tL(CAA)L, which was reported to contain a strong 5-dT termination signal (Braglia et al., 2005) (Figure S1A). Recruitment of Pol III to the promoter DNA and subsequent promoter DNA-opening was facilitated by pre-incubating the dsDNA with the Pol III GTF TFIIIB. To allow detection of the transcription products, the reaction mixture was supplemented with radioactively-labeled UTP. A time-course experiment showed the appearance of a strong RNA-band at the expected size of ca. 35 nucleotides (nt), which indicated that Pol III is capable to pause and presumably terminate on the 5-dT termination signal (Figure S1B). To confirm that the appearance of the RNA-band depended on the poly-dT termination signal, we tested and compared a 3-dT, 4-dT, and 5-dT oligo-dT sequence (Figure S1C). As expected, Pol III paused or terminated on the 5-dT signal but not on the sequences carrying only 3- or 4 dTs (Figure S1C). Instead, read-through (RT) products could be observed that corresponded to transcripts that were released once Pol III reached the end of the dsDNA. While our promoter-dependent transcription assay does not allow to distinguish between Pol III pausing and Pol III termination, a similarly prepared batch of purified Pol III could recently been shown to efficiently terminate on a 5-dT termination sequence on immobilized DNA templates (Xie et al., 2021). Together, these results confirmed that highly-purified Pol III is capable to pause, and presumably terminate, on its own *in vitro* on a poly-dT termination signal in a promoter-dependent transcription setup.

### A 7-dT signal traps Pol III in a pre-termination state

To determine the structure of the Pol III pre-termination complex (PTC), we first aimed to trap Pol III in the PTC conformation. To do so, we incubated yeast Pol III with a transcription scaffold that contained a 7-dT termination signal on the NT-DNA strand (Figure 1A). The scaffold also featured a 12-nt mismatch between the template (T)- and the NT-strand, thereby resembling a transcription bubble, in which the two strands are unwound, and the added RNA-oligonucleotide anneals to the T-DNA strand. We also substituted the DNA:RNA hybrid so that it doesn’t carry a poly-dA:rU sequence to increase the stability of the hybrid during scaffold preparation and to prevent spontaneous termination, which relies on the presence of a poly-dA:rU hybrid as an intrinsic destabilizing signal (Mishra and Maraia, 2019). To confirm that the 7-dT scaffold traps Pol III in a PTC-like state, we compared the capacity of Pol III to extend the RNA oligonucleotide in a 7-dT (PTC-like) or 2-dT (EC-like) scaffold (Figure 1B). As expected, addition of nucleoside triphosphates (NTPs) induced extension of the RNA primer for both scaffolds (Figure 1B). However, the activity of Pol III clearly differed between the two scaffolds. On the EC-scaffold, Pol III primarily transcribed long RNA products (lane 4). In sharp contrast, a strong RNA band, just three bases downstream of 3’-end of the used RNA primer was detected on the PTC-scaffold (lane 8). The appearance of this RNA band, which could hardly be observed on the EC-scaffold, indicated that Pol III pauses in the presence of the 7-dT signal on the NT-strand. Because the scaffold does not contain a poly-dA:rU hybrid and, therefore, should not trigger RNA release, we concluded that the 7-dT scaffold traps Pol III in a PTC-like state and proceeded with the structure determination of the Pol III PTC.

**Figure 1.**
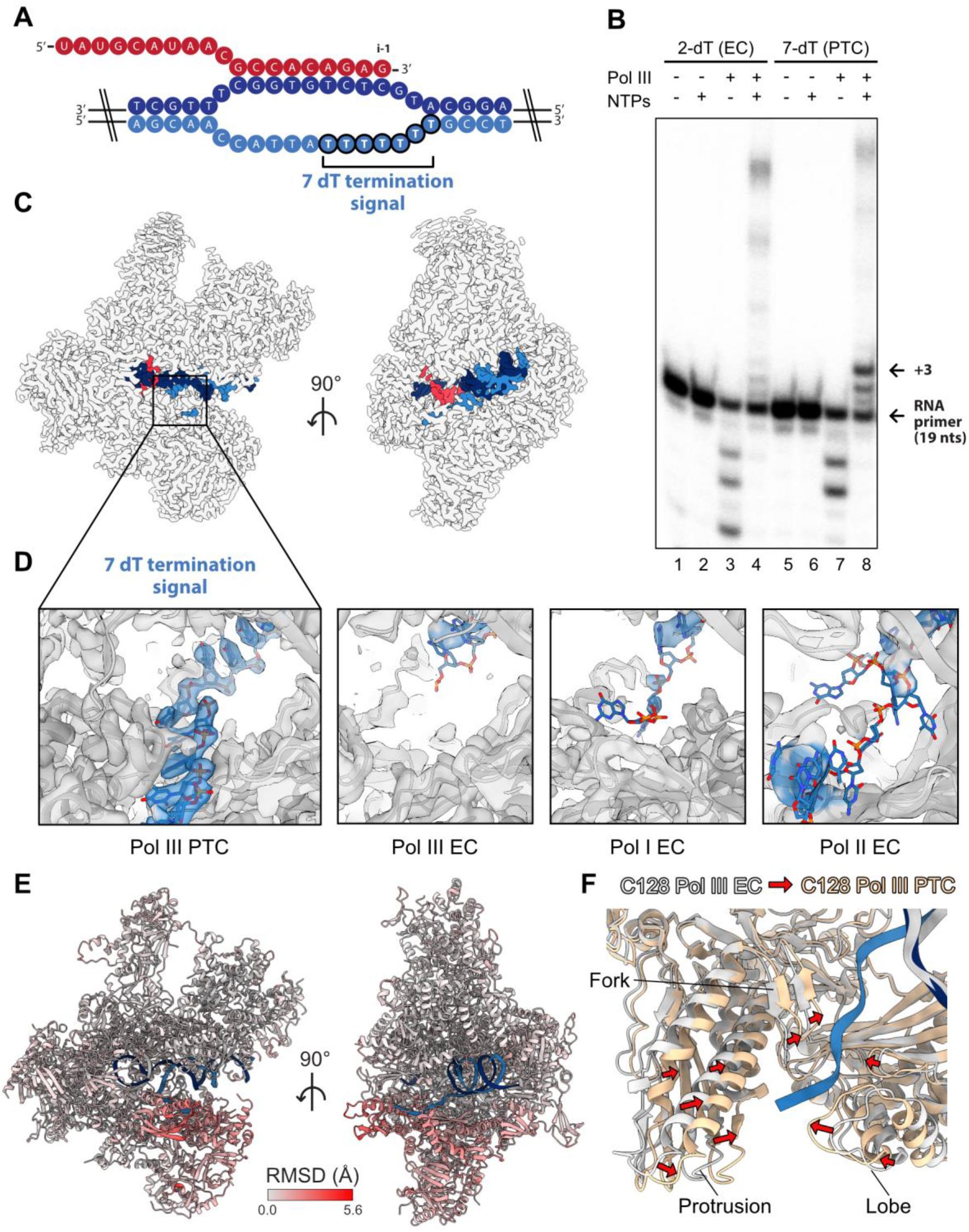
Structure of the yeast Pol III PTC. (A) Schematic of the 7-dT transcription scaffold. Red: RNA; dark blue: T-strand; light blue: NT-strand. The 7-dT termination signal is highlighted. The i-1 labels the position of the 3’-end of the RNA. (B) RNA primer extension experiment using a 5’ radioactively-labelled RNA oligonucleotide. Two scaffolds were tested: the elongation complex (EC) scaffold contains 2-dTs on the NT-strand (left), the PTC scaffold contains 7-dTs (right). n (Scaffold) = 2 pmol, n (Pol III) = 4 pmol. The RNAs were separated by a 15% TBE denaturing PAGE. (C) Cryo-EM reconstruction of the yeast Pol III PTC in two different views. Nucleic acids are colored as in (A). (D) Close-up view onto the NT-strand binding region of: Pol III PTC (this study), Pol III EC (this study), Pol I EC (EMDB: 0240, PDB: 6HLR), Pol II EC (PDB: 5C4J, 2Fobs-Fcalc calculated at 1.2 σ). (E) Atomic model of the yeast Pol III PTC colored by root-mean-square deviation (RMSD) between the Pol III EC (not shown) and Pol III PTC. (F) Close-up view illustrating the local contraction of subunit C128 around the NT-strand upon PTC formation from another view. C128 in the Pol III EC and PTC are colored in grey and beige, respectively.

### Structure of the yeast Pol III PTC

We assembled the yeast Pol III complex on the 7-dT PTC scaffold and acquired a high-resolution dataset on a Titan Krios transmission electron microscope (TEM), equipped with a K2 direct electron detector (Figure S2A). Processing the data yielded a 3D reconstruction of the Pol III PTC with a nominal resolution of 2.8 Å (Figure S2B) that locally extended to 2.5 Å in the Pol III core (Figure S2C). The cryo-EM map (map A) showed a clear signal for the bound nucleic acids with the downstream dsDNA and the DNA:RNA hybrid being well resolved (Figure 1C, Figure S3A). The C160 bridge helix and trigger loop, which are both important active site elements, were fully folded (Figure S3A). The C-terminal domain of C11, which was not resolved in the yeast Pol III EC structure, adopts an “outside funnel” conformation that is similar to the one in apo yeast Pol III (Hoffmann et al., 2015) but differs from the “outside funnel” conformation in human Pol III (Girbig et al., 2021) (Figure S3B). Thanks to the improved quality of the 3D reconstruction of the Pol III PTC, we could extend the Pol III structural model compared to existing models. Additional densities near the heterodimer and in the heterotrimer could be assigned to C53 (aa 195-224) and C31 (aa 101-161), respectively (Figure S3C, D). We further assigned additional densities, one in the DNA binding cleft formed by subunit C160 and one near subunit AC40 (aa 311-329) to molecules of the detergent CHAPSO, which was used to optimize cryo-EM grid preparation (Figure S3E, F).

Importantly, the cryo-EM reconstruction showed a strong signal of the unpaired NT-strand and allowed building of the complete 7-dT termination signal (Figure 1D). Because the cryo-EM sample preparation conditions differed between the Pol III PTC and the earlier reported Pol III EC (Hoffmann et al., 2015), we determined a control Pol III EC structure under exactly the same conditions as for the Pol III PTC and only changed the sequence of the NT-strand so that it contains only 2-dTs. High-resolution data collection and cryo-EM data processing (Figure S4A) yielded a 3D reconstruction of the Pol III EC control (map B) with a nominal resolution of 3.4 Å (Figure S4B) that locally reached 2.9 Å in the core (Figure S4C). While the cryo-EM signals of unrelated polypeptide elements were of comparable qualities, the NT-strand was only poorly resolved in the Pol III EC (Figure 1D). We further compared our cryo-EM reconstruction with reported structures of other eukaryotic RNA polymerases. The comparison revealed that the NT-strand in the Pol III PTC is also much better resolved than the NT-strand in the yeast Pol I EC (Tafur et al., 2016) and yeast Pol II EC structures (Barnes et al., 2015) (Figure 1D). Hence, the strong signal of the NT-strand is a specific feature of the Pol III PTC and indicates that Pol III is capable to recognize the poly-dT termination signal in a specific manner.

Superimposition with the Pol III EC control further shed light on conformational changes upon recognition of the termination signal as shown by a local contraction of subunit C128 around the NT-strand (Figure 1E, F). The contracting regions of C128 can be mapped to the C128 lobe (aa 196-370), protrusion (aa 60-195 and 371-439) and fork (aa 440-521) domains. In addition, the C53-C37 heterodimer, which binds the C128 protrusion domain, subunit C11, and the jaw domain of C160 contract towards the direction where NT-strand resides.

#### Subunit C128 specifically recognizes the termination signal

The high-resolution structure of the Pol III PTC enabled us to analyze in fine detail how Pol III recognizes the poly-dT termination signal. The NT-strand is exclusively bound by subunit C128 and is embedded into a positively-charged groove (Figure 2A). C128 forms 14 hydrogen bonds (H-bonds) with either the phosphate-backbone or the bases of the 7 dTs (Figure 2B). The residues that bind the NT-strand all reside in the lobe, protrusion and fork domains that contract upon the recognition of the termination signal (Figure 1 E, F). In addition, the methyl groups of dT_3_ and dT_4_ are embedded into a hydrophobic cavity formed by the fork loop 1 element (aa 440-453), thereby allowing Pol III to distinguish between a thymine and a cytosine base (Figure 2C).

Next, we analyzed to what extent the residues that bind the NT-strand are conserved and performed a multiple-sequence alignment (MSA) of C128 orthologs from 15 eukaryotic species covering a wide range of the eukaryotic tree of life (Keeling and Burki, 2019): ophistokonts (7), archaeplastids (3), stramenopiles (2), alveolata (2), and amoebozoans (1). Subjecting the MSA to the Scorecons software (Valdar, 2002) to retrieve conservation scores revealed a high degree of sequence conservation of the residues that form H-bonds with the NT-strand (Figure 2D). Four amino acids (Q199, K228, K393, R446) are identical in all analyzed super groups, and five amino acids have a high conservation score of 0.8-0.9 (S229, N245, R451, Q476, R481), which highlights the importance of these residues. Another four residues (K230, S246, K448, E450) exhibit moderate conservation scores of 0.5-0.7, and only one residue (T311) is not conserved across the eukaryotic tree. We also asked to what extent these residues are conserved across *S. cerevisiae* C128 and its two paralogs in Pol I (A135) and Pol II (RPB2). The MSA between the three paralogs revealed that 9 out of the 14 residues (Q199, K228, N245, S246, T311, K448, E450, Q476, R481) are unique to Pol III (Figure 2E). Hence, C128 utilizes a set of strongly conserved residues that are mostly unique to Pol III to engage tightly with the poly-dT termination signal.

**Figure 2.**
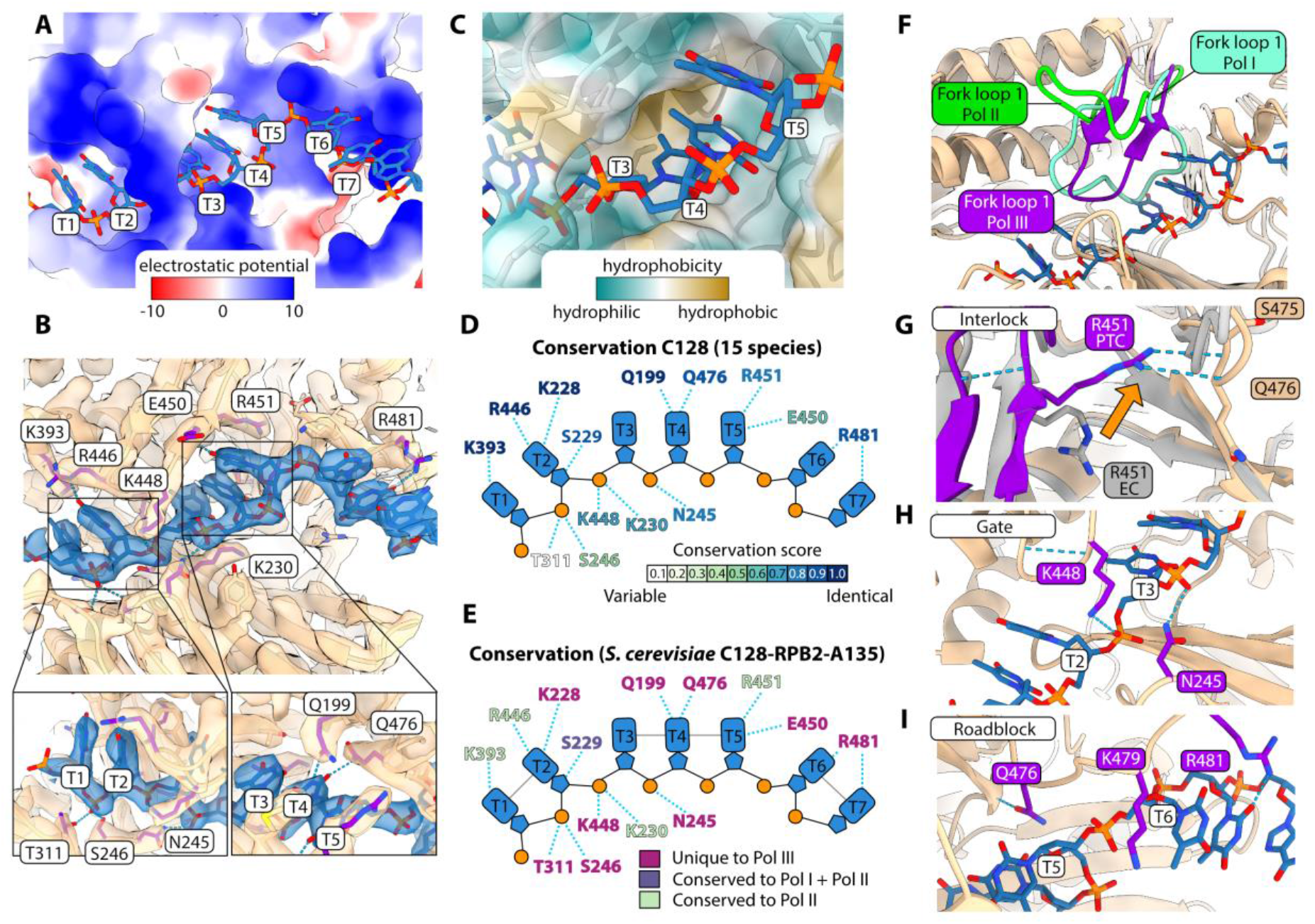
Termination signal recognition by C128 in the Pol III PTC. (A) Surface representation of C128, colored by electrostatic potential in kcal/(mol·e) at 298 K (red: negative, blue positive). The NT-strand, harboring the 7-dT termination signal is bound by a positively-charged groove. (B) Hydrogen-bond (H-bonds) between the 7-dT termination sequence (colored in blue) and C128 (colored in wheat). C128 residues that form H-bonds (blue dashed lines) with the NT-strand are labelled, and depicted as purple sticks. Shown are the atomic models and the corresponding cryo-EM densities as transparent surface. Bottom panels represent views from different angles. (C) Close-up view onto the hydrophobic groove formed by the fork loop 1 element of C128 that accommodates the methyl groups of dT_3_ and dT_4_. C128 is shown in the ribbon mode with its surface presentation in transparent and colored by hydrophobicity. (D) Nucleotide plot showing which residues of C128 form H-bonds with the NT-strand. The residues are colored according to their conservation scores that were derived from a multiple-sequence alignment (MSA) of C128 orthologs covering a diverse set of eukaryotic species. H-bonds are depicted as blue dashed lines. Conservation scores were calculated via Scorecons (Valdar, 2002). (E) Conservation between the H-bond contacting residues in C128 and its *S. cerevisiae* paralogs A135 (Pol I) and Rpb2 (Pol II). The residues are colored according to the legend below. (F) Fork loop 1 folds differently between Pol I, II, and III. The fork loop 1 of C128 (Pol III PTC, this study), A135 (Pol I EC; PDB: 6HLR), and Rpb2 (Pol II EC; PDB: 5C4J) are shown. (G) Close-up view on C128 residue R451 functioning as an interlock. The fork loops in the Pol III PTC and EC are colored in purple and grey, respectively. R451 changes its conformation upon EC to PTC transition as depicted by an orange arrow. (H) Close-up view on C128 residue K448 forming together with N245 a molecular gate that guides the NT-strand and breaks base-stacking between dT_2_ and dT_3_. (I) Close-up view on the fork loop 2. H-bonds between the NT-strand and Q476, R481 are shown. K479 defines the path of NT-strand and disrupts base-stacking between dT_5_ and dT_6_, thereby functioning as a molecular roadblock.

Multiple key residues that contribute to NT-strand recognition can be mapped to the fork loop 1 and fork loop 2 (aa 475-486) elements of subunit C128. Interestingly, fork loop 1 clearly folds differently in Pol III when comparing this region to the equivalent one in its counterparts Pol I and Pol II (Figure 2F). The fork loop 1 elements of Pol I (aa 469-483) and of Pol II (aa 466-478) adopt loop-like folds. In sharp contrast, fork loop 1 of Pol III folds into a β-hairpin, and its tip points into a different conformation compared to Pol II. This conformation is required to form the hydrophobic cavity of C128 and positions four residues (R446, K448, E450, R451) so that they can bind the NT-strand.

Superimposing the Pol III PTC and EC structures revealed that residue R451 in the fork loop 1 changes its conformation upon PTC formation (Figure 2G). In the Pol III EC, R451 points towards the coiled-coils of the C128 protrusion domain. During the EC to PTC transition, R451 flips by ca. 90° and forms two H-bonds with the peptide backbone of the fork loop 2 element (Figure 2G). We conclude that R451, thereby, functions as an interlock that keeps the fork loop 1 in a fixed position to define the path for the NT-strand. Another important residue is K448 that, together with the neighboring N245, forms a molecular gate that guides the NT-strand and disrupts the base-stacking between dT2 and dT_3_ (Figure 2H). Fork loop 2 bears three key residues for NT-strand coordination: Q476 forms an H-bond with the dT_4_ base, R481 forms two H-bonds with dT6 and dT7, respectively, and K479 functions as a positively charged molecular roadblock that serves as a spatial barrier for the NT-strand, interrupts the base stacking between dT_5_ and dT_6_ (Figure 2I), and guides the NT-strand to follow its defined path.

In summary, we interpret the tight H-bond network and the embedment of the methyl groups of the dT_3_ and dT_4_ bases as physicochemical barriers that counteract the translocation of Pol III along the DNA. Pol III uses a unique set of conserved residues, not only to form these physicochemical barriers but also to define the path of the NT-strand, which explains how termination signal recognition and pausing prior to termination is achieved.

### Mutating key dT-interacting residues in C128 interferes with transcription termination *in vitro*

To confirm the physiological relevance of our structural analysis of the Pol III PTC, we performed complementary functional experiments. We used CRISPR-Cas9 genome engineering of yeast cells to introduce several single-residue point mutations in subunit C128: Q199R, K448A, R451V, R481G, and H225L. In addition, we generated a C128 mutant that harbored three substitutions in the fork loop 2 region: Q476A, K479A, R481G (hereinafter referred to as Triple mutant). Q199 was chosen because it forms an H-bond with dT4, is unique to Pol III, and is conserved in all analyzed eukaryotic species. K448 forms the molecular gate that guides the NT-strand. R451 functions as an interlock keeping fork loop 1 in a fixed position. R481 forms two H-bonds with dT_5_ and dT6. H225 was chosen because it sits in the lobe region of C128 and forms a potential H-bond with D72 residing in a helix that contracts upon PTC formation. By mutating H225, we asked if other residues that do not contact the NT-strand but are, potentially, required to maintain the contracted state, also have a function in termination. The Triple mutant region contains substitutions in two H-bond forming residues (Q476, R481) and in K479, which acts as the molecular roadblock. In all residues, except one (Q476), we were guided by the MSA of Pol subunits C128, RPB2 and A135 to replace the C128 residues with their counterparts in RPB2 (Pol II-like) or A135 (Pol I-like). We, thereby, anticipated that the introduced mutations should not interfere with folding of the polymerase core elements.

For introducing the point mutations, we used a *S. cerevisiae* strain with a C-terminal TAP-tagged C128 subunit, which allowed us purifying endogenous WT and mutant Pol III variants to homogeneity for biochemical studies (Figure 3A). We next tested the effect of the C128 mutants using an *in vitro* transcription assay with a tailed-template containing a single-stranded overhang on the T-strand, to allow Pol III to initiate transcription independently of any transcription factors (Figure 3B). The constructs contained either a 6-dT termination signal or a 3-dT signal, through which Pol III should be able to transcribe without pausing or terminating. Furthermore, we designed the construct to contain the first guanine base (dG) on the NT-strand downstream of the termination sequence so that Pol III RT products can be captured by using only CTP, ATP and UPT in the transcription reaction. As expected, WT Pol III paused or terminated efficiently on the 6-dT termination signal as shown by a robust signal at the position where the termination signal sits (Figure 3C, lane 2). The 3-dT reaction, in contrast, did not show any sign of termination and, instead, RT products accumulated at the first dG (lane 11). Unexpectedly, we also observed additional RT bands upstream of the first dG. This could be due to a potential GTP contamination in the transcription reaction mix, which, in our point of view, does not affect the conclusions drawn below. Strikingly, we observed a strong reduction of the pausing or termination efficiency for all mutants, except for the H225L (Figure 3C, lane 3-8 and Figure 3D). We observed the strongest reduction in pausing/termination efficiency for the Q199R mutant (ca. 4-fold), followed by the Triple and R451V mutants (both ca. 3-fold). The K448A and R481G mutants showed a more modest, but still clear, reduction (ca. 1.7-fold). For the Q199R and R451V mutants, we also noticed an increase of RT products (Figure 3C, lane 4, 5). The tested mutants did not show any significant changes in the production of RT transcripts on the 3-dT control construct. Hence, the tested Pol III mutations only affected Pol III pausing or termination on the poly-dT sequence but not transcription *per se*. We would like to note that our experimental set-up did not allow distinguishing between defects in pausing or termination. Because we introduced the mutations to interfere with the recognition of the poly-dT signal, which serves as a pausing signal, it is likely that the observed effects actually reflect pausing defects. Noteworthy, the two processes are tightly coupled and pausing is a strict requirement for Pol III termination to take place (Arimbasseri and Maraia, 2013, 2015; Rijal and Maraia, 2016). Accordingly, any mutations that interfere with Pol III pausing will, most likely, also impair termination. The tailed-template transcription assay, thus, confirmed the importance of the amino acids Q199 (H-bond with dT4), R451 (interlocks fork loop 1), Q476 (H-bond with dT4), K479 (molecular roadblock) and R481 (H-bonds with dT_6_ and dT7). It also showed that the K448 molecular gate functions in transcription termination but it seems less important than the other tested residues as its substitution showed a less-severe pausing/termination deficiency. The H225L mutant, on the contrary, did not significantly interfere with termination.

**Figure 3.**
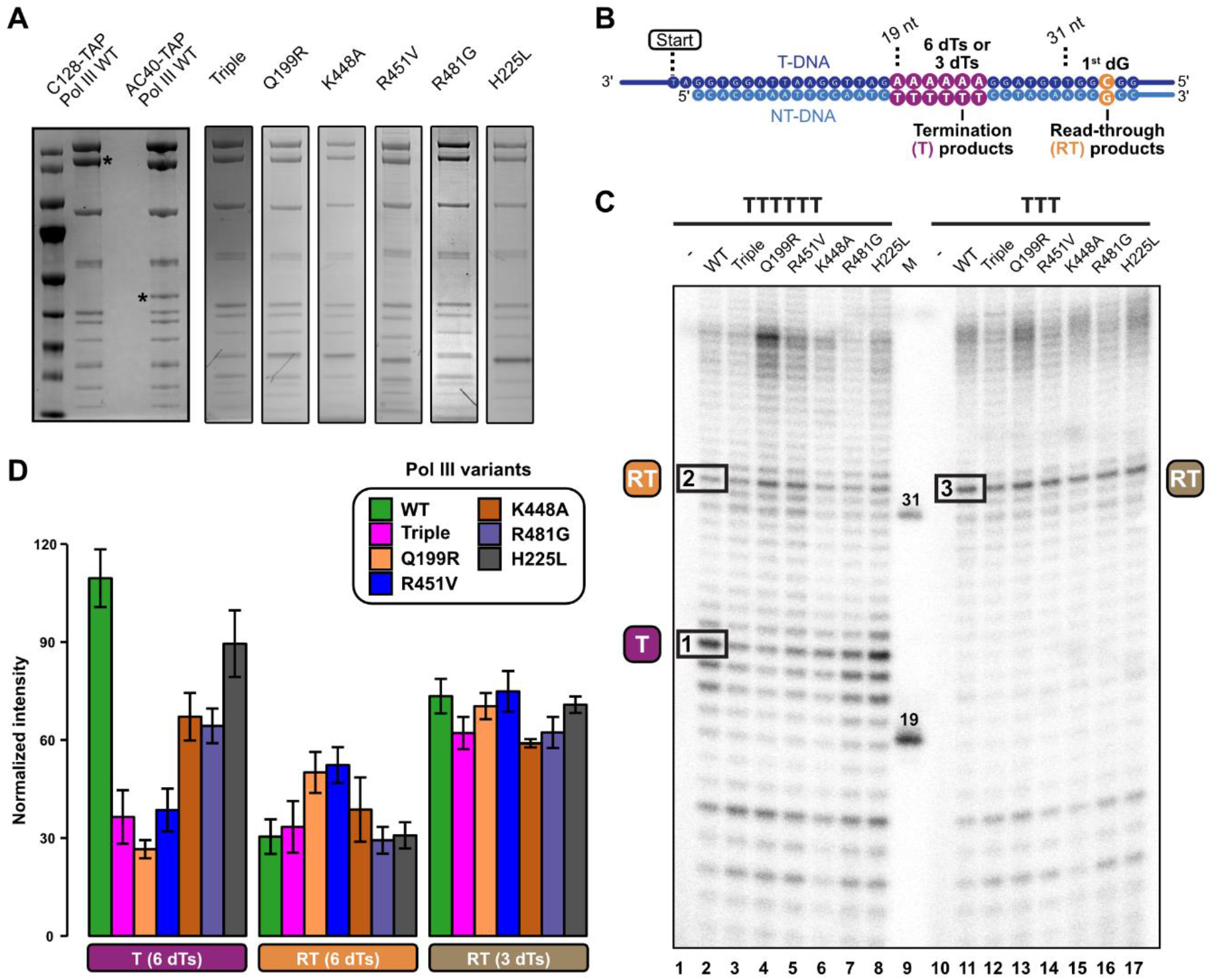
Mutating Pol III C128 on specific sites interferes with Pol III termination *in vitro*. (A) Protein gels of purified *S. cerevisiae* Pol III WT and mutant samples used for biochemical experiments. Asterisks mark the tagged subunits in the C128-TAP (used for the experiments) and AC40-TAP (loaded as control) Pol III samples. (B) Schematic of the tailed-template DNA construct. The positions of the poly-dT termination signal and of the first guanine base (dG) are highlighted. (C) *In vitro* transcription experiment on tailed-template DNA constructs. Experiments on WT and mutant Pol III variants were done on either a 6-dT or a 3-dT construct. The denaturing RNA gel corresponds to one out of three independent experiments. RNA bands used to quantify the Pol III transcription products are indicated by black boxes for the Pol III WT lanes. T - Termination; RT - Read-through. (D) Quantification of the Pol III transcription products indicated in panel (C). Bar plots represent normalized RNA band intensities (mean ± SD). For lane 15 (K448A), n = 2. RNA products were normalized by dividing the signal of the respective band by its total lane intensity, multiplied by a factor 1000.

### Pol III C128 mutations affect yeast viability and cause global termination defects *in vivo*

We next asked whether the C128 mutations also impact Pol III function *in vivo* and first compared the growth of the WT and the different C128 mutant strains (Figure 4A). The Triple mutation did not affect yeast growth at permissive temperature but caused a severe thermo-sensitive growth phenotype at 37 °C. We could also observe a slight thermo-sensitive phenotype for the R481G mutant. The Q199R displayed a clear growth defect at 30 °C, which was even more pronounced at 25 °C. These results support the idea that these mutations compromise Pol III function *in vivo*. For the K448A and R451V mutants, we did not observe any growth phenotype, suggesting that the effect of these mutations on Pol III activity is not severe enough to impair yeast growth.

We then directly assessed whether the most disruptive C128 mutations provoke Pol III termination defects *in vivo*. To this end, we generated high-resolution genome-wide maps of transcribing Pol III by CRAC (UV crosslinking and analysis of cDNAs), which allows mapping the position of Pol III with nucleotide resolution (Candelli et al., 2018; Granneman et al., 2009). We performed these experiments in strains carrying the WT, Q199R, R451V or the Triple mutant version of C128 (Figure 4 and Figure S5). We obtained very specific and reproducible Pol III signals in the two biological replicates, with most of the reads mapping to Pol III-transcribed genes (i.e. tRNA, 5S RNA and a few additional genes; Figure S5). Metagene analysis of Pol III distribution around tRNA genes (Figure 4B) indicates that a considerable fraction of wild type Pol IIIs reads through the primary terminator (i.e. the first T-tract after the 3’ end of the gene), in agreement with previous studies (Turowski et al., 2016; Xie et al., 2021). Importantly, we observed a significant increase in Pol III signal downstream of the primary terminator in the three mutants, indicating global transcription termination defects (Figure 4B and individual examples in Figure 4C). Heatmap analysis of Pol III occupancy (log2 ratio) in each mutant relative to the WT showed that such an increase in the Pol III RT signal can be detected for the vast majority of tRNA genes (Figure 4D), further confirming the generality of termination defects. Among these mutants, Q199R, which already showed a growth phenotype at permissive temperature, exhibited the most pronounced termination defects. While the R451V mutant did not show a growth phenotype, it was defective in Pol III transcription termination, similarly to the Triple mutant or even to a slightly more degree for some genes.

To perform a more quantitative assessment of termination defects caused by the different C128 mutations, we computed for each tRNA gene the RT index (determined by the ratio of Pol III signals within 500 bp downstream of the primary terminator relative to the gene body, Figure 4G) in the WT and mutant strains. Consistent with our metagene and heatmap analyses, we observed an increase in the average RT index, and thus a decrease in the Pol III termination efficiency, in all mutants compared to the WT, with the largest increase in Q199R and a more moderate and similar increase in R451V and the Triple mutant (Figure 4H). We considered the possibility that the different C128 mutations could impact differently the individual tRNA genes depending on the length of their terminator sequence. For instance, the Q199R mutation, predicted to prevent the interaction with the 4th T of the terminator, might not strongly affect termination at very long T-tracts. To address this possibility, we analyzed the RT index on tRNA genes grouped according the length of their primary terminator. However, we observed a similar behavior in the three mutants, namely a general increase in the RT index at all genes sets that tended to decrease as the length of the primary terminator increases (Figure 4I). Therefore, the introduced mutations reduce the efficiency of termination independently of the T-tract length.

Altogether, these results indicate that mutations in C128 designed to interfere with the recognition or the response to the poly-dT termination signal impair Pol III transcription termination *in vivo*.

**Figure 4.**
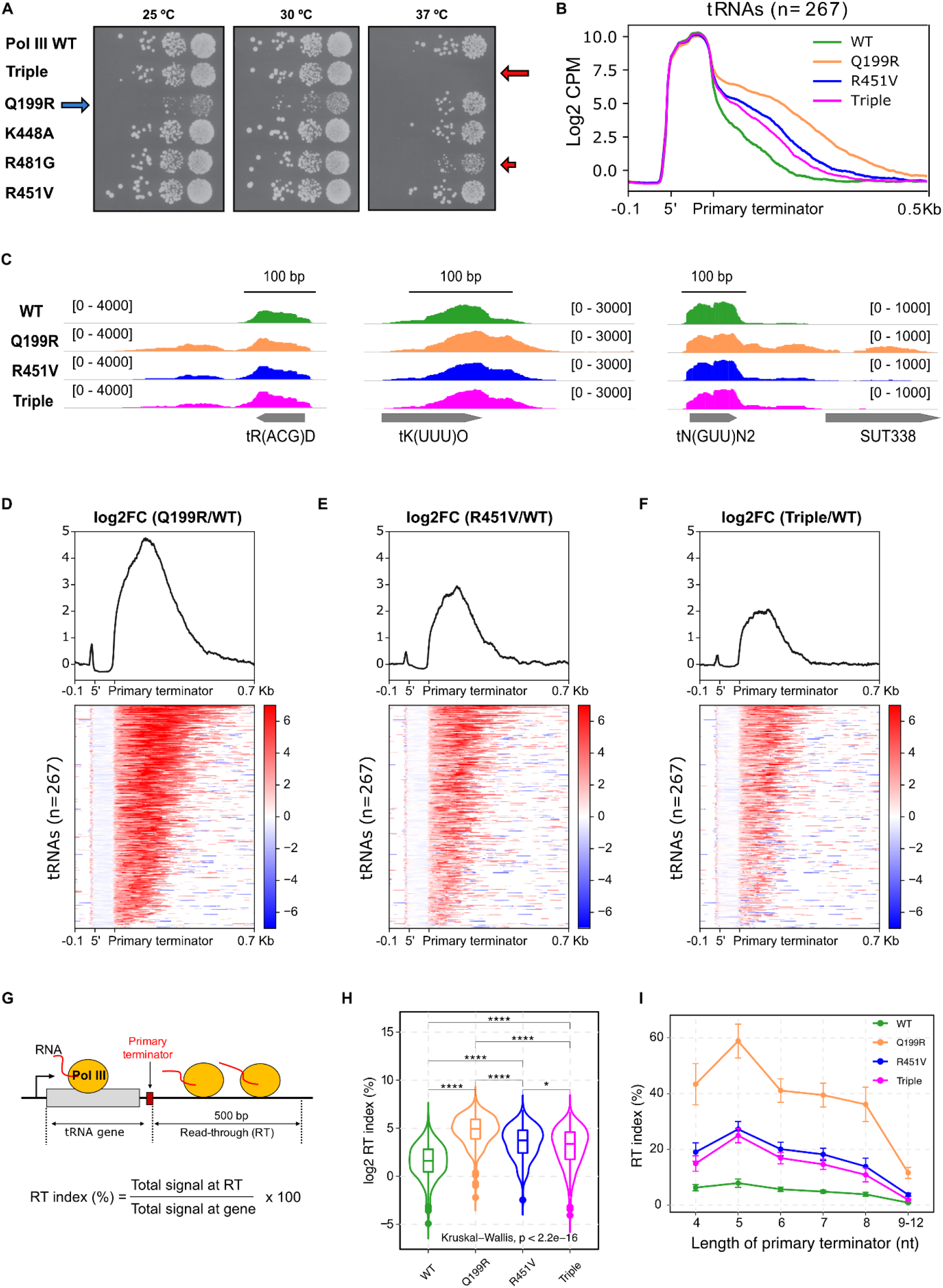
Characterization of the effect of C128 mutations on cell growth and Pol III transcription termination *in vivo*. **A)** Serial dilution analysis of the growth of the different C128 mutant strains at various temperatures. **B)** Metagene analysis of the Pol III distribution around tRNA genes. The signal covering the region between the 5’ end and the primary terminator (i.e. the 1^st^ T-tract after the 3’ end of the mature tRNA) is scaled to 100 bp. Values on the *y*-axis correspond to the mean coverage expressed as counts per million (CPM). **C)** Integrative Genomics Viewer (IGV) screenshots of examples of tRNA genes displaying termination defects in the indicated strains. The values under brackets correspond to the scale of the RNAPIII signal expressed in CPM. Only the signal on the strand hosting the tRNA gene is indicated. **D-F)** Heatmap analyses representing the log2 fold change (FC) of the Pol III signal in the indicated mutants relative to the WT at tRNA genes. The summary plot on the top was calculated using the average values for each position. Note that tRNA genes in the three heatmaps are sorted in the same order determined by the ranking of signals in D. **G)** Scheme of a tRNA transcription unit indicating the gene body and the read-through (RT) region used for calculating the RT index. **H)** Comparison of the average RT index for tRNA genes in the indicated strains. *p* corresponds to the p-value for the global comparison of the four groups according to the Kruskal-Wallis test. Asterisks denote the p-values of pairwise comparisons (*: p ≤ 0.05; **: p ≤ 0.01; ***: p ≤ 0.001; ****: p ≤ 0.0001). **I)** Analysis of the RT index of tRNA genes grouped according to the T-track length of their primary terminator in the indicated strains. Data points correspond to the average value whereas error bars denote the standard error.

### Structure of Pol III Δ sheds light on the role of C53-C37 in Pol III transcription termination

The structure of the Pol III PTC and the associated functional studies revealed a key role of subunit C128 in Pol III transcription termination. However, they did not explain the function of the C53-C37 heterodimer, which has been demonstrated to be essential for Pol III transcription termination (Arimbasseri and Maraia, 2013, 2015; Landrieux et al., 2006; Rijal and Maraia, 2013). C53-C37 binds the lobe domain of C128 and, thereby, is positioned relatively close to the NT-strand. However, C53-C37 does not bind the NT-strand directly, raising the question of how it functions in transcription termination. To better understand the role of C53-C37, we determined the structure of a Pol III variant, which lacks the C53-C37 heterodimer and subunit C11 (hereinafter referred to as Pol III Δ). We purified Pol III Δ from a *S. cerevisiae* strain, in which the endogenous C11 gene was substituted with its *S. pombe* ortholog (Kassavetis et al., 2010). The *S. pombe* substitution permits yeast viability but causes loss of C53-C37 and C11 during the protein purification procedure (Kassavetis et al., 2010). The Pol III Δ complex was purified to homogeneity (Figure 5A), assembled on the 7-dT PTC transcription scaffold, and subjected to cryo-EM analysis (Figure S6A). The 3D reconstruction (map C) had a nominal resolution of 3.9 Å (Figure S6B) that locally reached 3.7 Å in the Pol III core (Figure S5C). Despite the lower resolution compared to the Pol III PTC, the cryo-EM map was still of sufficient quality to build the Pol III Δ structure for comparison with the Pol III PTC structure. As expected, the Pol III Δ structure lacked the density for C53-C37 and C11 but clearly featured cryo-EM density for the bound DNA and RNA elements (Figure 5B). Interestingly though, the cryo-EM signal corresponding to the NT-strand was much poorer than the one derived from the Pol III PTC structure, despite the presence of the 7-dT termination signal (Figure 5B). We only observed fragmented density for the NT-strand at lower threshold levels, which was not of sufficient quality to confidently build the model of the unwound NT-strand. When we superimposed the Pol III PTC and Pol III Δ structures, we further noticed that the fork, protrusion and lobe elements of C128 adopted a relaxed, instead of a contracted conformation (Figure 5B, C). Hence, the earlier described contraction of the Pol III core upon NT-strand recognition was essentially reversed in the absence of C53-C37 and C11. Instead, the conformation of the Pol III Δ core resembled that of the Pol III EC despite the presence of the termination signal (Figure 5D). We concluded from this observation that the C53-C37 heterodimer – although it does not directly contact the NT-strand – indirectly assists Pol III termination by binding the lobe of C128 and positioning it for interaction with the NT strand.

**Figure 5.**
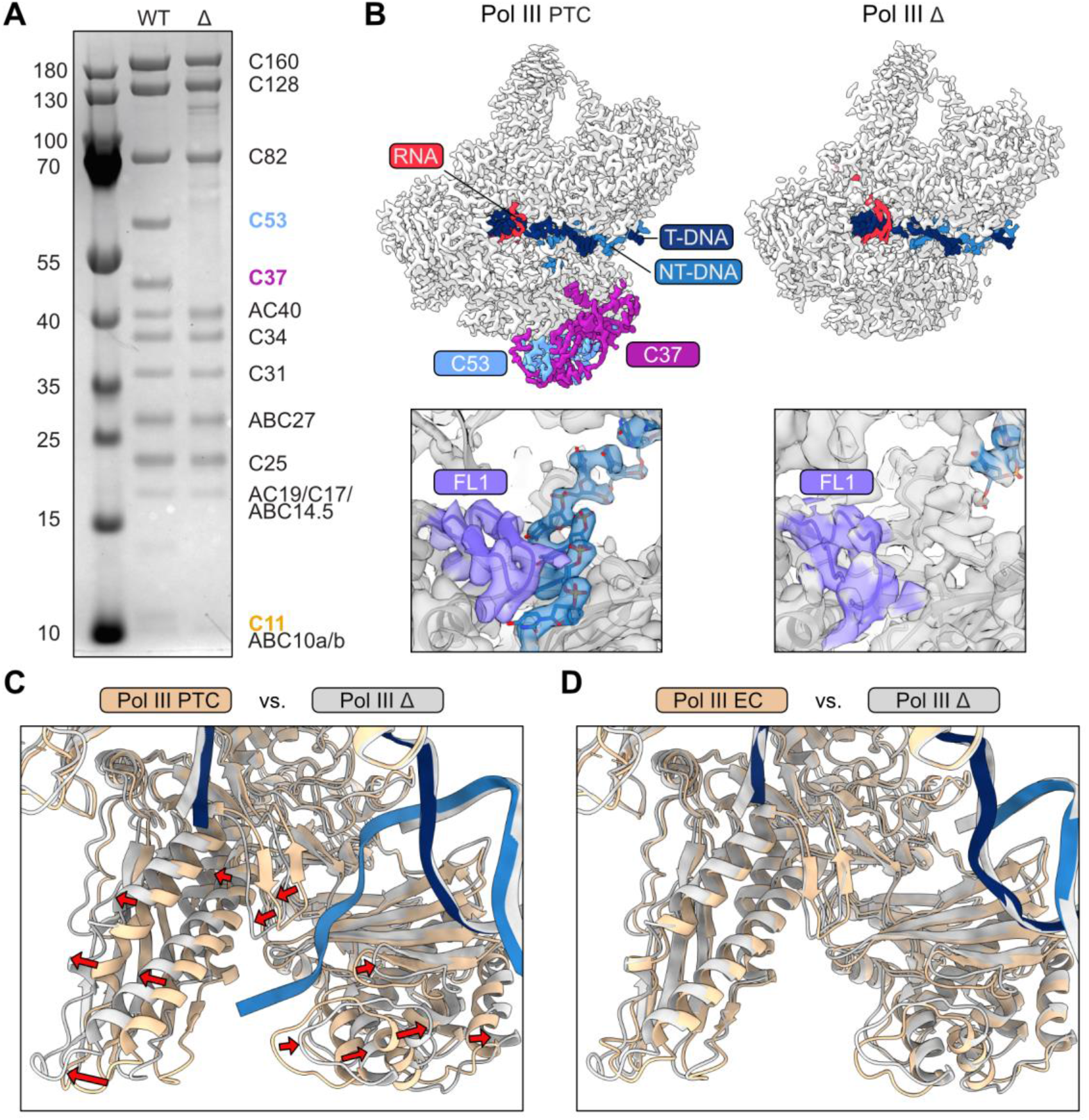
The structure of Pol III Δ gives insights into the role of C53-C37 in NT-strand recognition. (A) Protein gel of Pol III WT and Pol III Δ, which lacks C53-C37-C11. The names of the subunits absent in Pol III Δ are shown in color. Parts of this gel image (marker and WT lanes) have already been used for a figure as part of another study (Xie et al., 2021). (B) Top: Comparison between the cryo-EM maps of the Pol III PTC (left) and Pol III Δ (right). The Pol III Δ map lacks the C53-C37 heterodimer that is highlighted in the Pol III PTC map. Bottom: close-up views onto the NT-strand binding region in the Pol III PTC. Cryo-EM densities are shown as transparent surfaces. The NT-strand is poorly resolved in the Pol III Δ map. Fork loop 1 (FL1) is colored in purple. (C) Superimposition of the Pol III and Δ Pol III PTC structures. Conformational changes are indicated by red arrows. (D) Superimposition of the Pol III and Δ Pol III EC structures.

### Structure of Pol III PTC with nucleotides reveals a register offset between the unwound DNA strands

Finally, to obtain another snapshot of the Pol III termination mechanism, we set out to determine the structure of the Pol III PTC in the presence of NTPs. We incubated Pol III with the 7-dT PTC scaffold and with an NTP mix containing CTP, ATP, and UTP, collected a high-resolution cryo-EM data set and obtained 3D reconstruction (map D) of the Pol III PTC + NTPs (Figure 6A) at a nominal resolution of 2.7 Å that locally extended to 2.5 Å (Figure S7A-C). When processing the cryo-EM data set, we noticed that the upstream DNA was slightly better resolved, which encouraged us to perform masked 3D classification on the nucleic acid elements (Figure S6D, E). We, thereby, obtained a 3D map (map E) at 3.2 Å that featured a significantly improved signal of the upstream DNA moiety (Figure 6B). The 3D reconstruction enabled us to recognize the DNA minor groove and allows the visualization of the complete transcription bubble (Figure 6B).

Importantly, the cryo-EM densities in the Pol III PTC and PTC + NTP structures were of sufficient qualities to exactly define the registers of the DNA:RNA hybrids in the active sites (Figure 6C, D). In the Pol III PTC, the 3’-end guanine (G19) of the used RNA oligonucleotide sits on position i-1, whereas the nucleotide addition side (i+1) position is not occupied (Figure 6C). In the Pol III PTC + NTPs structure, the RNA was extended by at least one nucleotide, and the bases of the hybrid were shifted by one position away from the nucleotide addition site (Figure 6D). The G19 base was, consequently, placed on position i-2, and the i-1 site was occupied by a newly attached cytosine (C20). We also observed a weak density that occupied position i+1 and extended the RNA model with another adenosine (A21). Surprisingly though, we did not observe a register shift on the NT-strand (Figure 6D, E), which was also very well resolved in the Pol III PTC + NTPs structure. Hence, the addition of NTPs induced a register shift of the DNA:RNA hybrid but left the NT-strand unchanged. We speculate that the induced register offset between the two DNA strands traps Pol III in an unproductive state as discussed below in more detail.

**Figure 6.**
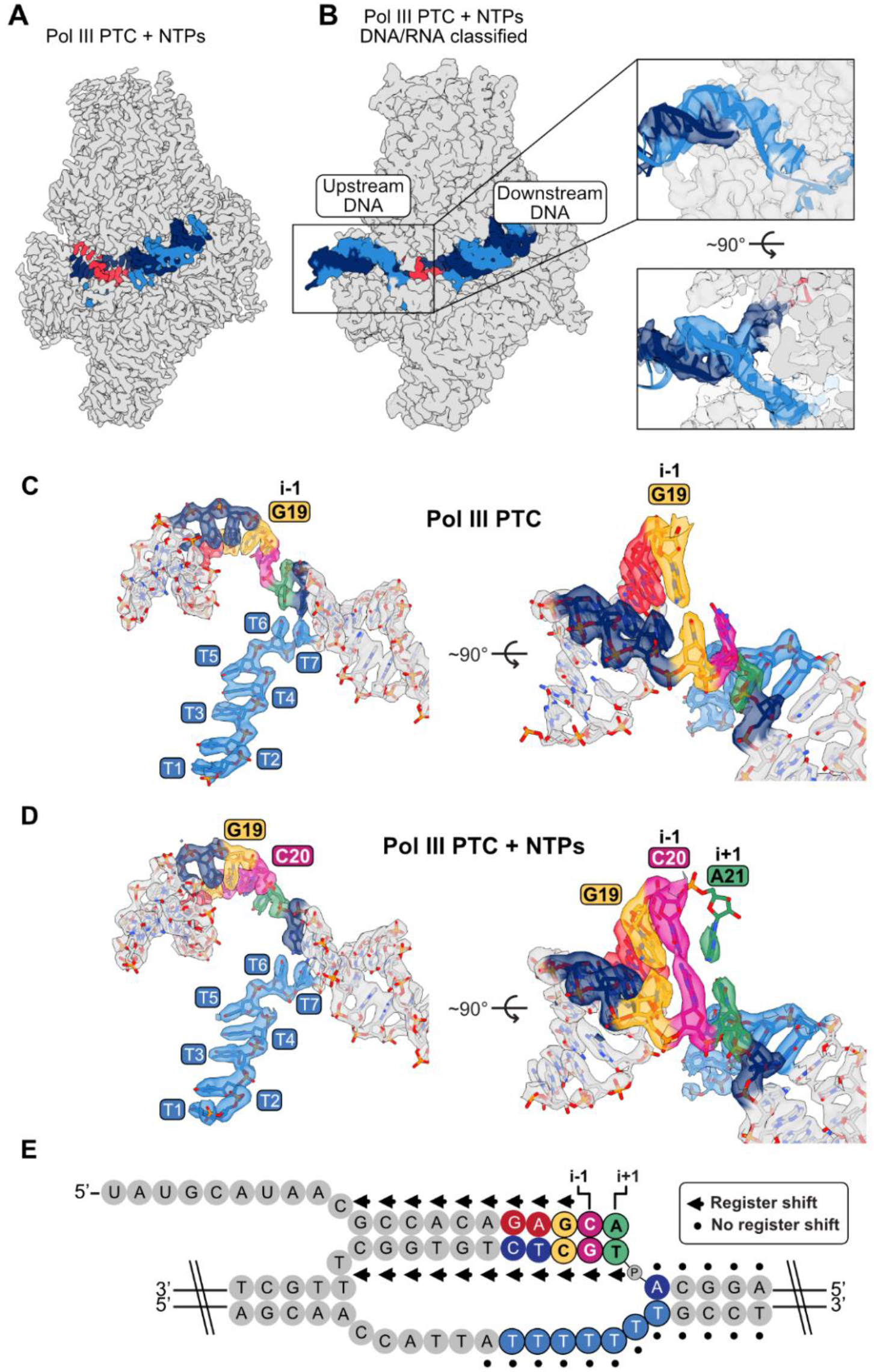
Structure of the Pol III PTC + NTPs reveals a register-offset in the Pol III PTC upon addition of NTPs. (A) Cryo-EM map of Pol III PTC + NTPs. Nucleic acids are colored in dark blue (T-DNA), light blue (NT-DNA), and red (RNA). (B) Cryo-EM map of Pol III PTC + NTPs with improved signal for the upstream DNA, derived via masked classification on nucleic acid elements. A putative nucleic acid strand model for the upstream DNA is shown for illustrative purposes but was not included into final Pol III PTC+NTPs model because of too much uncertainty about the correct path of the DNA backbone. (C) Close-up views on the transcription bubble in the Pol III PTC. Nucleotides of interest are colored to illustrate the register of the DNA:RNA hybrid and of the NT-strand in the Pol III PTC. The G19 base at position i-1, resembling the 3’-end of the RNA, is colored in orange. (D) Close-up views on the transcription bubble in the Pol III PTC + NTPs structure. Nucleotides are colored as in (C) to illustrate RNA-strand extension upon addition of NTPs. The i-1 position is occupied by C_20_ (pink) that has been added by the Pol III. The nucleotide addition site (i+1) also shows weak density (green) for another added nucleotide. The register of NT-strand did not change between Pol III PTC and Pol III PTC + NTPs. (E) Schematic of the poly-dT transcription bubble highlighting the observed register shift (arrows) in the DNA:RNA hybrid but not on the NT-strand (dots) upon addition of NTPs. Thereby, a register offset is generated that may trap Pol III in an unproductive state. Nucleotides are colored as in (C) and (D).

## DISCUSSION

### Model of Pol III transcription termination

Terminating on poly-dT termination signals is a unique feature of the Pol III transcription machinery and is a crucial specialization of Pol III to cope with the high demand of its transcripts. Here, we determined the high-resolution cryo-EM structure of the yeast Pol III PTC at a resolution of 2.8 Å, which revealed in fine detail how Pol III (but not Pol I and Pol II) is able to recognize and pause on the poly-dT termination signal. Using a combination of genetic, biochemical, and high-resolution genomic approaches, we have thoroughly validated the conclusions of our structural analyses both *in vitro* and *in vivo*. We also determined complementary structures of the Pol III ΔC53-C37-C11 variant and of the Pol III PTC in the presence of NTPs. Combining our structural and functional investigation of the Pol III PTC with the insights obtained by us and others in previously published work, allows us to propose a mechanistic model for Pol III transcription termination and to describe how this process might be integrated into the transcription cycle (Figure 7).

In the first step of the transcription cycle, Pol III initiates transcription with the help of general transcription factors (Step 1). Once the complete transcription bubble forms, Pol III escapes the promoter and enters the elongation phase. In the Pol III EC, the NT-strand only loosely associates with the core of Pol III and, therefore, is too flexible to be captured structurally. When Pol III reaches a termination site that harbors at least 4 dT bases, the C128 subunit recognizes the termination signal, Pol III pauses and the Pol III PTC forms (Step 2). PTC formation is driven by a tight H-bond network between subunit C128 and the termination sequence. The NT-strand becomes ordered, and the Pol III core contracts around it. Fork loop 1 establishes multiple H-bonds with the NT-strand and forms a hydrophobic cavity that accommodates the methyl groups of two dT bases. C128-mediated recognition of the termination signal is aided by the C53-C37 heterodimer, which binds the C128 lobe domain. We earlier hypothesized that a flexible loop in C37 (aa 197-224), which connects two helices that bind the C128 lobe domain, might participate in Pol III transcription termination by directly binding the NT-strand (Hoffmann et al., 2015). Given that we did not observe a direct interaction between this loop and the termination signal in the Pol III PTC structure, we need to revise this hypothesis. We propose that binding of C53-C37 reduces the flexibility of C128 and, thereby, reinforces the tight engagement of C128 with the NT-strand when the poly-dT sequence is exposed. In the absence of C53-C37, Pol III would not be slowed down once it has reached the termination signal, thus preventing correct termination. This mode of function rationalizes the well-known role of C53-C37 in transcription termination, which was, so far, only poorly understood on a molecular level.

The tight interaction between the NT-strand and C128 induces pausing of Pol III. Nevertheless, the polymerase still appears to be able to extend the RNA molecule, which “pulls” the T-strand towards the active site (Step 3). The strong interaction with the termination sequence, however, hinders further translocation along the DNA and induces a register offset between the T- and the NT-strand. Such offset could trap Pol III in an unproductive state that might be sensed by the TFIIS-like RNA cleaving subunit C11 (Step 4). C11 has been demonstrated biochemically to be required for Pol III transcription termination (Chedin et al., 1998; Landrieux et al., 2006; Mishra and Maraia, 2019; Mishra et al., 2021). C11 is homologous to TFIIS, which is needed in the Pol II system to reactivate stalled Pol II by cleaving off backtracked RNA (Cheung and Cramer, 2011). We, therefore, speculate that C11 also can sense an unproductive state of Pol III.

In attempts to rescue the stalled Pol III PTC, C11 may insert its catalytic C-terminal domain into the Pol III funnel. Insertion of C11 into the Pol III funnel has been recently shown to be concomitant to partial opening of the Pol III clamp domain that contacts the downstream DNA (Girbig et al., 2021) (Step 5). Because transcription termination does not dependent on the RNA-cleavage activity of C11 (Landrieux et al., 2006; Mishra and Maraia, 2019; Mishra et al., 2021), we propose that C11 insertion into the Pol III funnel serves as a destabilizing signal by opening the clamp domain so that Pol III loses its “grip” on the downstream DNA. Owing to the poly-dT termination signal on the NT-strand, a poly-adenine:uracil (dA:rU) hybrid forms in the active site, which is much less stable than hybrids with other sequences (Martin and Tinoco, 1980) (Step 6). The funnel insertion of C11 could reinforce dismantling of the already unstable hybrid. Since the DNA:RNA hybrid itself is an important stabilizing element of the Pol III EC, the falling-apart of the hybrid would consequently cause the Pol III PTC to collapse and to release both the DNA and the nascent RNA molecule (Step 7).

**Figure 7.**
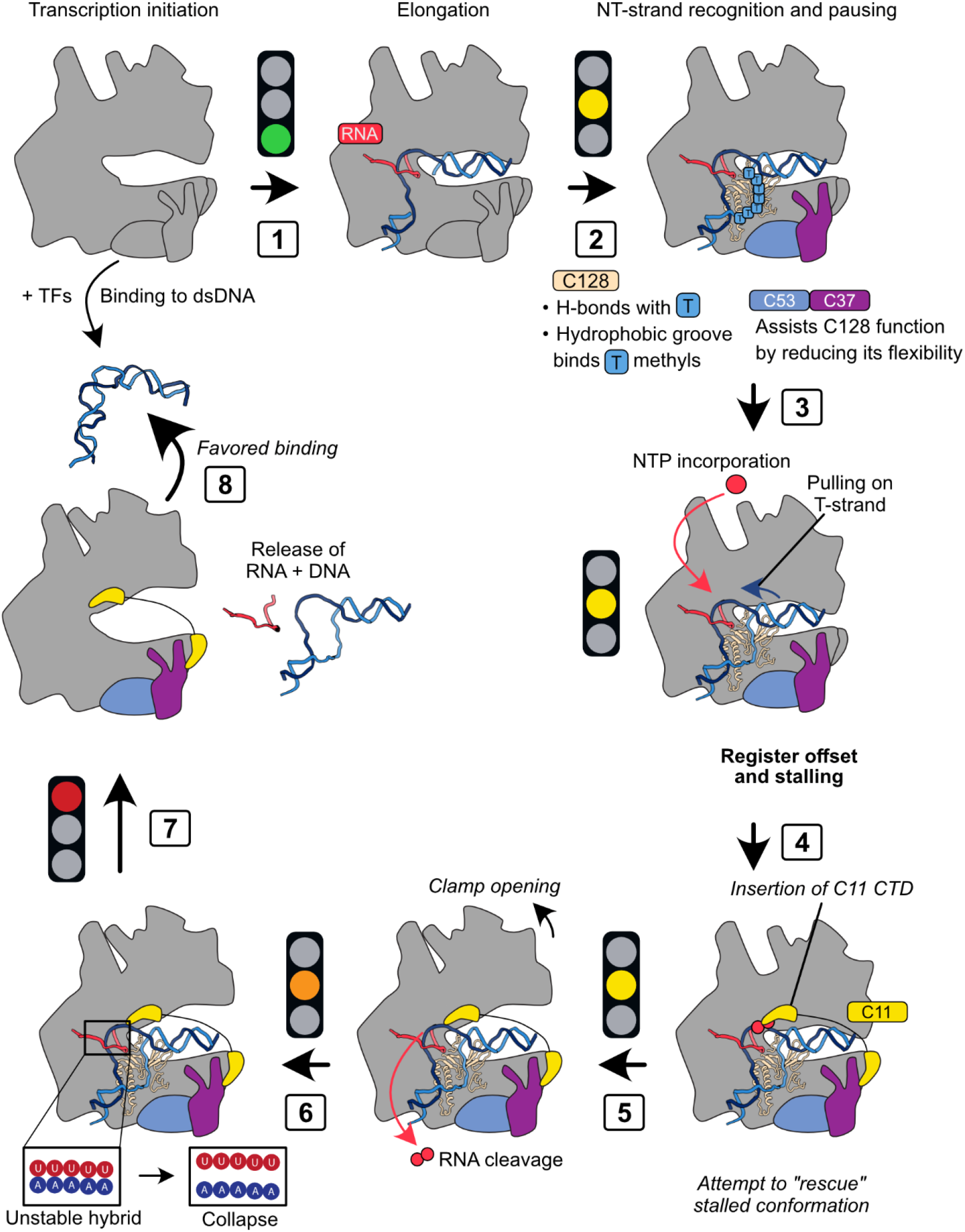
Model of Pol III transcription termination. The model combines the here-presented structural and biochemical data with literature-based hypotheses (italicized).

C11 is not only required for transcription termination but also for facilitated recycling of Pol III (Dieci and Sentenac, 1996; Landrieux et al., 2006). Only recently, this mode of function could be assigned to the linker domain that links the N-terminal and the C-terminal domain (CTD) of C11 (Mishra and Maraia, 2019). We speculate that a terminated Pol III that still harbors the C11 CTD inserted into its funnel has an increased propensity to reinitiate transcription (Step 8). The cryo-EM structures of the yeast Pol III initiation complexes revealed that the clamp adopts a closed conformation in the Pol III PIC (Abascal-Palacios et al., 2018; Vorländer et al., 2018) and, thus, needs to first open up to allow recruitment of Pol III to the promoter DNA. We propose that this process could be assisted by the insertion of C11 into the funnel.

### Comparison between the yeast and human Pol III PTC

When this manuscript was in preparation, the structure of the human Pol III PTC at a resolution of 3.6 Å was reported (Hou et al., 2021). The human Pol III PTC structure showed comparable characteristics to our yeast PTC structure. The six bases of the poly-dT termination signal were well resolved and multiple H-bonds formed between the termination signal and RPC2 (the human C128 ortholog). The authors also reported a local contraction of the lobe, protrusion and fork elements around the NT-strand, which agrees well with our findings. Together with our biochemical investigation of the key residues in C128, our investigation of complementary structures (Pol III Δ and Pol III PTC + NTPs), and the in vivo validation of its physiological relevance, the two Pol III PTC structures support and complement each other and together significantly advance our mechanistic understanding of the Pol III termination process.

Notably, the number of the dTs that are required for Pol III transcription termination differs between human and yeast. Whereas *S. cerevisiae* Pol III needs at least 5 dTs to terminate, the majority of human tRNA genes only possess a 4-dT termination site (Braglia et al., 2005), which is why human Pol III must be more sensible to shorter dT-tracts. Given that the human and yeast Pol III PTC appear to recognize the NT-strand in a similar manner, we speculate that the differences between the two species is either due to a less stably bound DNA:RNA hybrid in the active site or to differences in the contribution of the C11 subunit (RPC10 in humans) to termination. Indeed, we noticed that the DNA:RNA hybrid in the human Pol III PTC was less well resolved than in the yeast Pol III PTC. However, we could not observe any obvious differences in the amino acids that contact the DNA:RNA hybrids in the two structures that might account for the observed differences. We therefore suggest that the observed differences pertain to the impact of C11/RPC10. In both the yeast and human Pol III PTC structures, the C-terminal domains of C11/RPC10 are positioned outside of the Pol III funnel (but they adopt different conformations). As we reported recently, human RPC10 is able to dynamically monitor the active site of Pol III (Girbig et al., 2021), which became apparent by the fact that RPC10 either adopts an “outside funnel” (Girbig et al., 2021) or “inside funnel” conformation (Girbig et al., 2021; Li et al., 2021; Wang et al., 2021). A similar conformation, in which the CTD of C11 penetrates the Pol III active site, has not yet been captured for yeast Pol III, indicating that human RPC10 has a higher tendency to enter the Pol III active site. We hypothesize that the active site of human Pol III is, owing to the action of human RPC10, more sensitive to unstable poly-dA:rU hybrids than yeast Pol III, which could explain why human Pol III requires less dTs to terminate transcription.

## Acknowledgements

We thank F. Weis and W.J.H. Hagen (EMBL Cryo-Electron Microscopy Service Platform) for EM support, T. Hoffmann and J. Pecar and EMBL IT Support for computational and data storage support and T. Hoffmann and J. Pecar in particular for setting-up and maintaining the high-performance computing environment, S.C. Vonesch for advice on CRISPR-Cas9-mediated genome engineering of yeast cells, F. Baudin for RNA labeling and advice on biochemical experiments, M.K. Vorländer for providing purified TFIIIB components, and advice on cryo-EM data collection- and processing, N.A. Hoffman for initial project discussions, and current members of the Müller lab for discussions. M.G., H.G. and C.W.M acknowledge support by EMBL. M.G. was supported by a Boehringer Ingelheim Fonds PhD fellowship. D.L. and O.P. were supported by the Agence National pour la Recherche (ANR-16-CE12-0001-01 to O.P. and ANR-16-CE12-0022-01 to D.L) and the Fondation pour la Recherche Medical (F.R.M., programme Equipes 2019). J.X. was supported by the China Scholarship Council, by the FRM (FDT202012010433) and the LabEx “Who Am I?” (ANR-11-LABX-0071 and the Université de Paris IdEx ANR-18-IDEX-0001) funded by the French Government through its “Investments for the Future”.

## Author contributions

C.W.M. initiated and supervised the project. M.G. carried out protein purifications, biochemical experiments, cryo-EM grid preparation and data collections, model building and interpreted the structures. H.G. performed genome engineering of yeast cells via CRISPR-Cas9, handled yeast cultures, and did yeast spotting assays. J.X. performed CRAC experiments. J.X., D.L., and O.P. analyzed and interpreted CRAC results. M.G. wrote the manuscript with input from the other authors.

## METHODS

### Large scale purification of endogenous Pol III

Yeast Pol III was purified in large scale from a *S. cerevisiae* strain (Euroscarf ID #SC1613) carrying a C-terminal TAP-tag on subunit AC40. Large-scale purified Pol III was used for structural analysis and biochemical experiments. Yeast cells were grown in a 100 L fermenter (BIOSTAT® D-DCU, Sartorius) in YPAD medium for 17 h at 30 °C. Cells were harvested in log phase (OD_600 nm_ > 6.5) with a flow-through centrifuge, which typically yielded a ca. 1.5 kg cell pellet that was kept at -80 °C until usage. For purification, the yeast cell pellet was thawed over night at 4 °C and, the next day, washed with cell wash buffer (50 mM Tris-HCl pH 8 (RT), 250 mM (NH_4_)_2_SO_4_, 12 mM β-mercaptoethanol) by dissolving the yeast pellet in 100 mL wash buffer per 100 g cell pellet and centrifugation (JLA 8.1000 rotor, Beckman) at 5,000 rpm, 4 °C for 10 min. The supernatant was removed and the cell pellet was resuspended in lysis buffer (250 mM Tris-HCl pH 8 (RT), 20% glycerol, 250 mM (NH_4_)_2_SO_4_, 1 mM EDTA, 10 mM MgCl_2_, 10 μM ZnCl_2_, 12 mM mercaptoethanol), supplemented with protease inhibitor (PI) cocktail mix (0.3 ug/mL Leupeptin, 1.4 ug/mL Pepstatin, 170 μg/mL PMSF, 330 μg/mL benzamidin, in EtOH). For cell lysis, all material and equipment was pre-cooled at 4 °C and lysis steps were performed at 4 °C. Cells were lysed with glass beads in a bead beater (BioSpec) using ca. 230 mL cell suspension per beaker (ca. 6 beakers in total) and 170 mL glass beads per beaker, supplemented with a spatula tip of Pefabloc. Beakers were cooled using an ice-salty water (5 M NaCl) bath. Cells were lysed by beating 8 times for 45 s and waiting for 90 s. The lysate was filtered through a metal mesh and centrifuged (JA-14 rotor, Beckman) at 14,000 rpm, 4 °C for 1 h. The supernatant was recovered, filtered again through a metal mesh and applied to a 350 mL heparin sepharose (GE Healthcare) column equilibrated in HEP bind buffer (50 mM Tris-HCl pH 8 (RT), 20% glycerol, 250 mM (NH_4_)_2_SO_4_, 0.5 mM EDTA, 1 mM MgCl2, 10 μM ZnCl_2_, 1 mM mercaptoethanol), supplemented with PI mix (see above). The remaining cell pellet (containing unlysed yeast cells) was resuspended in lysis buffer, lysed again using the procedure described above, and also applied to the heparin column. Bound proteins were eluted using a 400 mL linear gradient (0-100%) from buffer HEP A (50 mM Tris-HCl pH 8 (RT), 250 mM (NH_4_)_2_SO_4_, 0.5 mM EDTA, 1 mM MgCl_2_, 10 μM ZnCl_2_, 1 mM mercaptoethanol, PI mix) into HEP B (50 mM Tris-HCl pH 8 (RT), 1 M (NH_4_)_2_SO_4_, 0.5 mM EDTA, 1 mM MgCl_2_, 10 μM ZnCl_2_, 1 mM mercaptoethanol, 0.5 mM PMSF). The eluate was diluted with HEP A buffer to 500 mM (NH_4_)_2_SO_4_ and incubated with 20 mL IgG sepharose beads (GE Healthcare), equilibrated by washing with first 0.5 M AcOH pH 3.4 and second 1M Tris-HCl pH 7.5 (RT) (repeated twice) 4 °C, rolling over night. The next day, beads were washed with 200 mL IgG binding buffer (50 mM Tris–HCl pH 8, 10% glycerol, 225 mM (NH_4_)_2_SO_4_, 0.5 mM EDTA, 1 mM MgCl_2_, 10 µM ZnCl_2_, 2 mM β-mercaptoethanol), and bound proteins were eluted via addition of Tobacco etch virus (TEV) protease, incubation for 6 h at 4 °C, and recovery of the supernatant. 100 mL of IgG elution buffer (50 mM Tris–HCl pH 8, 60 mM (NH_4_)_2_SO_4_, 0.5 mM EDTA, 1 mM MgCl_2_, 10 µM ZnCl_2_, 2 mM β-mercaptoethanol) were added to the IgG beads and the flow-through was recovered and pooled with the first eluate. The eluted protein solution was applied to a MonoQ 10/100 column (Sigma-Aldrich, Merck), equilibrated in MonoQ buffer A (40 mM Hepes, pH 7.5, 60 mM (NH_4_)_2_SO_4_, 0.5 mM EDTA, 1 mM MgCl_2_, 10 µM ZnCl_2_, 10 mM DTT). Bound proteins were eluted via linear gradient from 60 to 1000 mM (NH_4_)_2_SO_4_ in buffer and buffer exchanged into EM buffer (15 mM Hepes, pH 7.5, 150 mM (NH_4_)_2_SO_4_, 10 mM DTT). The protein sample was concentrated to 7.5 -14.9 mg/mL, snap-frozen in liquid nitrogen and stored at -80 °C until usage.

### Small scale purification of endogenous Pol III

Yeast Pol III was purified in small scale for structure-function studies from either a *S. cerevisiae* strain (Euroscarf ID #SC2330) carrying a C-terminal TAP-tag on subunit C128 (Pol III WT) or from *S. cerevisiae* mutant strains, genome-engineered via CRISPR-Cas9 (see below). Yeast cells were grown in 24 L cultures (in YPD medium) in conical flasks. To do so, 30 mL of yeast over-night cultures were prepared and diluted to 100 mL the next morning. In the evening, the cultures were diluted to 500 mL. The next day, the cultures were diluted to an OD_600 nm_ of 0.2, split amongst 16 x 1.5 L cultures and grown to an OD_600 nm_ of 1.5. The cultures were harvested by centrifugation (JLA 8.1000 rotor, Beckman, 3000 rpm, 10 min), washed once with PBS, and snap frozen in liquid nitrogen. Frozen cells (ca. 50 g per 24 L) were thawed over night at 4 °C and resuspended in lysis buffer (see above), supplemented with PI mix (see above). The cells were lysed as described above. The lysate was cleared via centrifugation and loaded on two 5 mL HiTrap Heparin HP (Cytiva, Merck) columns (connected in series) that were equilibrated in lysis buffer. Proteins were eluted using a 100 mL linear gradient (0.25 to 1 M (NH_4_)_2_SO_4_). Pol III was further purified as described above except that the MonoQ step was omitted. Buffer-exchanged proteins were concentrated up to 0.4 to 5.6 mg/mL, snap frozen, and stored at -80 °C.

### Purification of yeast Pol III Δ

Pol III Δ (lacking C53/C37/C11) was purified for structural analysis from an *S. cerevisiae* strain harboring a His6-tag on subunit C128. The strain was a gift from Richard Maraia (National Institutes of Health, Bethesda,MD, USA) and features a substitution of endogenous *S. cerevisiae* C11 with *S. pombe* C11, which causes loss of the C53/C37 heterodimer upon protein purification (Kassavetis et al., 2010). Yeast cells were grown in a 100 L fermenter culture in YPGAL medium to an OD_600 nm_ of ca. 5.0-6.0. A 330 g pellet of frozen cells was thawed over night at 4 °C and resuspended in lysis buffer (60 mM Hepes pH 7.5, 750 mM NaCl, 10.5 mM MgCl_2_, 7.5% glycerol, 14.3 mM mercaptoethanol), supplemented with PI mix (see above). The cells were lysed as described above. The lysate was cleared via two centrifugation runs (1 h, 4 °C, each): first run at 30,000 g; second run at 72,465 g. The lysate was loaded overnight onto a 5mL HisTrap HP column (Cytiva, Merck) equilibrated in 20 mM Hepes pH 7.5, 500 mM NaCl, 7 mM MgCl_2_, 10% glycerol, 10 mM mercaptoethanol, 10 mM imidazole). Bound proteins were eluted using a 100 mL linear gradient (10 to 300 mM imidazole), diluted with 120 mL of MonoQ buffer (40 mM Tris-HCl pH 8, 68 mM (NH_4_)_2_SO_4_, 0.5 mM EDTA, 1 mM MgCl2, 10 μM ZnCl2, 10 mM DTT) and loaded onto a MonoQ 5/50 column (Sigma-Aldrich, Merck). Bound proteins were eluted using a 30 CV linear gradient (68 mM to 1 M (NH_4_)_2_SO_4_), diluted with MonoQ A buffer, and applied a second time to the Mono Q 5/50 GL to increase purity. For the second elution step, a two-step linear gradient (first step: 68 mM to 250 mM (NH_4_)_2_SO_4_; second step: 250 mM to 1 M (NH_4_)_2_SO_4_) was used. Eluted proteins were buffer exchanged as described above, concentrated to 2.4 mg/mL, snap-frozen in liquid nitrogen and stored at -80°C until usage.

### Preparation of transcription scaffolds

Transcription scaffolds were always prepared freshly before usage. Scaffolds used for cryo-EM sample preparation were annealed in a thermocycler (Bio-Rad) in the following way: oligonucleotides were dissolved in water. 1 μL of the template-strand and 1 μL of the non-template strand (100 μM each) were mixed and incubated at 95 °C for 3 min. The temperature was incrementally reduced to 25 °C at a rate of 1 °C/min. 3 μL of 2x hybridization buffer (2xHB) (40 mM Hepes pH 7.5, 24 mM MgCl_2_, 200 mM NaCl, 20 mM DTT) were added. 1 μL of the RNA strand (100 μM) were pre-heated at 45 °C for 3 min and added to the mixture, which was incubated at 45 °C for 3 min. The temperature was reduced to 20 °C at a rate of 0.7 °C/min, and the annealed scaffold was kept on ice.

### Cryo-EM sample preparation

All samples were diluted to the working concentrations using EM buffer (15 mM Hepes, pH 7.5, 150 mM (NH_4_)_2_SO_4_, 5 mM MgCl2, 10 mM DTT), supplemented with 4 mM (final concentration) CHAPSO. For assembly of the nucleic-acid bound complexes, Pol III protein samples were incubated at 4 °C for 60 min with 1.05x molar excess of pre-annealed transcription scaffolds (see above). The following final protein concentrations were used: 2.5 μM (Pol III PTC); 2.8 μM (Pol III EC); 3.4 μM (Pol III PTC + NTPs), 2.6 μM (Pol III Δ). For the Pol III PTC + NTPs sample, the EM buffer was further supplemented with CTP, ATP, and UTP (10 mM each) and the Pol III-nucleic acid complex was incubated for 90-instead of 60 min. After incubation with transcription scaffolds, 3 μL of sample was applied to 200-mesh copper R2/1 grids (Quantifoil), which were glow-discharged twice using a Pelco EasyGlow instrument. The cryo-EM grids were prepared at 100% humidity and 4 °C using a Vitrobot Mark IV (Thermo Fisher Scientific) and the following blotting parameters: wait time 10 s, blot force 4, blot time 4 s. For cryo-EM sample preparation of Pol III Δ, a slightly different protocol was used: the DNA:RNA hybrid was first pre-annealed in a thermocycler by mixing 1 μL of T-DNA and 1 μL of RNA (100 μM each) using the same program as for the other annealing steps. After annealing, the DNA:RNA hybrid was diluted to a concentration of 12 μM in HB buffer (20 mM Hepes pH 7.5, 12 mM MgCl_2_, 100 mM NaCl, 10 mM DTT). Next, 9 μL of Pol III Δ protein sample (c = 4 μM) were mixed with 3 μL DNA:RNA hybrid (c = 12 μM), and the mixture was incubated at 28 °C for 10 min. Afterwards, 1 μL of NT-DNA (c = 40 μM, diluted in HB buffer) strand was added, and the reaction was incubated for another 10 min at 28 °C before supplementing the mixture with 4 mM CHAPSO (final concentration) and plunge freezing as described above.

### Cryo-EM data collection and processing

SerialEM (Mastronarde, 2005) was used for automated data collection. Cryo-EM data were collected on a Titan Krios transmission electron microscope (TEM) operated at 300 keV (Thermo Fisher Scientific), equipped with a Quantum energy filter (Gatan), and with a K2 (for Pol III PTC, EC and PTC + NTPs) or K3 (for Pol III Δ) direct detector (Gatan). Cryo-EM data collection parameters (magnification, pixel size, number of collected movies, movie frames, exposure time, dose, defocus range) for the four datasets are specified in Table S1. Warp (Tegunov and Cramer, 2019) was used for initial frame-alignment, dose-weighting, estimation of the contrast transfer function (CTF) and particle picking. The data was further processed in RELION 3.1 (Zivanov et al., 2018). For particle polishing, frame-aligned and dose-weighted was also done via RELION’s own implementation of the MotionCor2-algorithm (Zheng et al., 2017) and CTF-corrected with Gctf (Zhang, 2016). CryoSPARC (Punjani et al., 2017) was also used at earlier stages of image processing to assess data quality via 2D classification and generation of a 3D ab-initio reconstruction of the Pol III PTC, which was later used as a reference for 3D classification in RELION. Particle numbers and data processing steps are outlined in Supplementary Figs. S2, S3, S5, S6. In case of the Pol III PTC + NTPs dataset, the region corresponding to the upstream DNA was better resolved than in the other 3D reconstructions. We, therefore, performed masked 3D classification on the bound nucleic acids, which yielded a map that featured a clearly visible upstream DNA moiety (map E). Reported nominal resolution values of 3D reconstructions were calculated via RELION post-processing using the Fourier shell correlation (FSC) 0.143 cut-off criterion (Rosenthal and Henderson, 2003). Local resolution ranges were estimated using RELION.

### Structural model building and refinement

Cryo-EM maps, used for structural model building, were sharpened via Local Deblur (Ramírez-Aportela et al., 2019), implemented into the Scipion software framework (de la Rosa-Trevín et al., 2016). Scipion, in turn, was run via the SBGrid software collection (Morin et al., 2013). Structural model building was performed using Coot (Casañal et al., 2020; Emsley et al., 2010). Starting models were placed into cryo-EM density maps using UCSF Chimera (Goddard et al., 2007). The first map used for structural model building was of the Pol III PTC. The model of the yeast Pol III EC (Hoffmann et al., 2015) was used as the starting model (PDB: 5FJ8). The model was first real-space refined in Coot using distance restraints to accommodate for conformational changes of the newly build model. Distance restraints were computed using ProSMART (Nicholls et al., 2012), implemented into Coot. The sequence of the DNA:RNA scaffold was mutated to resemble the scaffold used in the Pol III PTC sample. Clear additional density could be observed of the NT-strand, and the sequence of the poly-dT termination signal was manually build into this density. The model was further manually adjusted without distance restraints to fit the cryo-EM density of the Pol III PTC and real-space refined in Coot using Ramachandran restraints. The Pol III PTC model was further subjected to ISOLDE (Croll, 2018) to reduce the clash score and optimize the model geometry. For the Pol III EC structure, the Pol III PTC model was used as a starting model. Conformational changes were first adjusted by subjecting the Pol III PTC model and the Pol III EC map to the Namdinator software (Kidmose et al., 2019). The sequence of the nucleic acids was adjusted to resemble the used EC scaffold. No density of the unwound NT-strand could be observed, and the corresponding region was deleted. The Pol III EC model was further manually inspected and adjusted in Coot. For the Pol III Δ model, the procedure was the same as for the Pol III EC except that the C53/C37 heterodimer and C11 were deleted from the Pol III PTC starting model. For the Pol PTC + NTPs structure, a register shift of the DNA:RNA hybrid could be observed when inspecting the density in the active site, and the DNA:RNA hybrid was adjusted accordingly. Weak density for a nucleotide base at the position i+1 was also noticed and the 3’-end of the RNA was, therefore, extended with an adenosine monophosphate molecule. Interpretation of the upstream DNA in map E was further aided by subjecting map E to LocScale (Jakobi et al., 2017). All models were refined multiple times against the corresponding cryo-EM maps with Phenix (Liebschner et al., 2019) using Ramachandran- and secondary structure restraints. ChimeraX (Goddard et al., 2018) was used to visualize cryo-EM maps and atomic models for the structural analysis and interpretation and for figure preparation.

### Primer extension experiments

For assembly of scaffolds used for primer extension experiments, the DNA and RNA oligonucleotides (1 μL, 20 μM each) were mixed and incubated at 95 °C for 3 min. The mixture was placed immediately on ice for 1 min. Afterwards, 3 μL of 2x HB buffer were added, and the mixture was incubated at room temperature for at least 10 minutes. Prior to scaffold assembly, the used RNA was radioactively labelled on the 5’-end with [γ-32P]ATP (10 mCi/ml, Hartmann Analytic) and T4 PNK (NEB), following purification via PAGE. The annealed scaffold was diluted to 2 μM with transcription reaction buffer (TRB) (20 mM Hepes, pH 7.6, 60 mM (NH_4_)_2_SO_4_, 10 mM MgSO4, 10% glycerol, 2 mM DTT). 1 μL of diluted scaffold was incubated with 1 μL of Pol III (4 μM, also diluted in TRB) for at least 10 min at RT. Primer extension was initiated via addition of 7 μL TRB, supplemented with NTPs (1 mM each, final concentration). The reaction was incubated at 28 °C for 20 min and stopped by adding 3 μL of formamide and boiling for 4 min. The reaction was subjected to PAGE using a denaturing (8M Urea) 17% polyacrylamide TBE gel. Radioactive signal was captured with a phosphor-imaging screen (Fujifilm) and detected with a Typhon FLA9500 (GE Healthcare) instrument.

### *In vitro* transcription experiments

Promoter-dependent and independent were performed to analyse Pol III transcription termination. Annealing of the DNA oligonucleotides (Sigma-Aldrich, desalt grade) was done similarly for the two types of experiments: 1 μL of template DNA-strand (30 μM) and 1 μL of non-template DNA-strand (30 μM) were mixed, boiled for 2 min, and immediately placed on ice for 1 min. 2 μL of 2x HB (see above) were added, and the mixture was incubated at room temperature for 10 min. The annealed DNA was diluted to a concentration of 2 μM using TRB (see above) and kept on ice until usage. For promoter-independent experiments, tailed-template DNA constructs were designed to carry a single-stranded DNA overhang on the template strand. In a 5 μL reaction volume, 2 pmol of DNA were mixed with 2.5 pmol of WT or mutant Pol III, which were pre-diluted to working concentrations (0.6 – 2.5 μM) using EM buffer. The mixture was incubated at room temperature for 10 min, and transcription was initiated by adding 5.3 μL of a transcription initiation mix (3.9 μL of TRB, 0.7 μL NTP mix (10 mM ATP, 10 mM CTP, 1 mM UTP) and 0.7 μL of [α-P32]UTP (10 mCi/ml)). The reactions were incubated at room temperature for 40 min. Afterwards, 138 μL of stop buffer (0.6 M NaAcetate, 30 mM EDTA, 0.5% EDTA) were added, and the reaction was subjected to phenol-chloroform extraction, boiling for 3 min, and denaturing PAGE using a 15% polyacrylamide TBE-Urea gel. Radioactive signal was recorded as described above. Band intensities were quantified via the Image Lab software (Bio-Rad). Plots were generated in R. To normalize the radioactive signals of the bands of interest, their band intensities were divided by total lane intensities that contained the respective band of interest. The calculated values were multiplied by a factor of thousand for illustrative purposes. Tailed-template experiments were performed in triplicates, except for the mutant K448A on the 3-dT construct, which was done in duplicates. For promoter-dependent transcription assays, a double-stranded (ds) DNA hybrid construct was designed that consisted of the *S. cerevisiae* U6 snRNA promoter region and a portion of the *S. cerevisiae* tDNA gene tL(CAA)L featuring a 5-dT termination signal. Nucleic acids were annealed as described above. Pol III was diluted to 5 μM using TRB. Yeast TFIIIB, which was used to recruit Pol III to the promoter region, was freshly assembled by mixing a fusion construct of Brf1 and TBP (Brf1-TBP) with Bdp1, which were expressed and purified as described in (Vorländer et al., 2018). TFIIIB was assembled by mixing 2.4 μL of Bdp1 (6 mg/mL), 2.8 μL of Brf1-TBP (6 mg/mL) and 4.8 μL of TRB-300 (20 mM Hepes, pH 7.6, 300 mM NaCl, 60 mM (NH_4_)_2_SO_4_, 10 mM MgSO4, 10% glycerol, 2 mM DTT). The mixture was incubated on ice for 10 min and diluted to 5 μM using TRB-300. Next, 5 pmol of assembled TBIIIB were incubated with 2.5 pmol of dsDNA on ice for 10 min. Subsequently, 5 pmol of Pol III were added and the mixture was incubated for at least 10 min on ice. Transcription was initiated by adding 5.4 μL of transcription initiation mix (3.9 μL of TRB, 0.7 μL NTP mix (10 mM ATP, 10 mM CTP, 10 mM GTP, 1 mM UTP) and 0.7 μL of [α-P32]UTP (10 mCi/ml)). The reaction was incubated at 28 °C. Typically, time courses were performed by stopping the reaction at the desired time points with 4 μL formamide loading buffer and boiling at 3 min. The reactions were directly loaded on 15% denaturing PAGE and radioactive transcription products were recorded as described above. The here-shown gels derived from promoter-dependent transcription experiments were not performed in triplicates but the reproducibility could be confirmed by performing the experiments multiple times for different analytical purposes.

### Mutagenesis of *S. cerevisiae* C128 for structure-function studies

CRISPR-Cas9-mediated genome engineering was used for site-directed mutagenesis of the endogenous *S. cerevisiae* C128-coding gene. A C128-TAP-tagged yeast strain (Euroscarf ID: SC2330) was used for genome engineering. Guide RNAs (sgRNA) were designed using the Benchling CRISPR Guide RNA Design tool (2021), retrieved from https://benchling.com, and ordered as DNA oligonucleotides in forward and reverse directions. For cloning of the sgRNA constructs, the oligonucleotides were phosphorylated via T4 PNK (NEB), mixed, and annealed by incubating the mixture at 95 °C for 6 min and letting it cooling down to room temperature. Next, 2 μL of the annealed dsDNA was mixed with 0.5 μL of the pWS082 entry vector (which was a gift from Tom Ellis lab and Addgene plasmid # 90516) (at a concentration of 200-300 ng/uL), 1 μL each of T4 ligase, 10x T4 ligase buffer, and BmsbI. In a thermocycler, the mixture was incubated at 42 °C (2 min) and 16 °C (5 min), and the cycle was repeated 10 times. Next, the reaction was incubated at 60 °C for 10 min and lastly at 80 °C for 10 min. The reaction was transformed into *E. coli* XL-1 Blue chemically competent cells (Agilent). The next day, positive clones were selected by using a Typhoon imager with a GFP-filter to screen for GFP-negative colonies. DNA donor constructs (containing the mutated DNA sequence to generate the corresponding C128 mutation) were designed to contain 50 base pairs (bps) homology arms at either side of the desired DNA break. The oligonucleotides were ordered as two partially complementary (20 bps) primers in forward and reverse direction, amplified via polymerase chain reaction (PCR), and purified via ethanol precipitation. The Cas9-sgRNA gap repair vector pWS174 (which was a gift from Tom Ellis lab and Addgene plasmid # 90961) was linearized with BsmBI (NEB) and gel-purified. The cloned sgRNA entry vectors were digested with EcoRV. For transformation into yeast cells, 5 ug of dsDNA donor DNA, 100 ng of linearized pWS174, and 200 ng of linearized sgRNA entry vector were mixed and transformed into yeast cells via the lithium acetate (LiAc)/sorbitol method. 50 mL of yeast cells at an OD_600 nm_ of 0.6 were spun down (2500 rpm, 5 min RT), washed once with a mix of LiAc (1M)/Sorbitol (1M) and resuspended in 600 μL LiAC/Sorbitol. The DNA mixture (above) was added to 100 μL of the yeast LiAc/Sorbitol suspension, and the mixture was added to a pre-mix containing 280 μL of 50% PEG 4000 and 15 μL sperm carrier DNA. The reaction was incubated at RT for 1 hour. 40 μL of DMSO were added, and the reaction was incubated at 42 °C for 20 min while slowly shaking. Next, 1 mL of YPD were added, followed by centrifugation (3000 rpm, 2 min). The pellet was dissolved in 500 μL YPD, transferred into 15 mL tubes containing 4.5 mL YPD, and the mixture was incubated at 30 °C for a least 3 hours. The cell suspension was spun at 2300 rpm for 5 min, dissolved in 500 μL PBS, spread on YPD plates supplemented with Nourseothricin (clonNAT) (100 ug/mL), and kept at 30 °C. After three days, yeast colonies were re-streaked on YPD plates without clonNAT and incubated for another two or three days. Positive clones (those who contain the desired mutation) were screened via colony PCR and subsequent Sanger sequencing.

### Yeast spotting assays

YPD medium was inoculated with yeast cultures, and the yeast starter cultures were grown over night at 30 °C shaking. The next day, the OD_600 nm_ was measured, the cultures were diluted to an OD_600 nm_ of 0.2 using fresh YPD, and the cultures were incubated for another 4 hours. Next, the cultures were diluted to OD_600 nm_ = 0.2 using PBS. The PBD-diluted cell suspensions were used to make 10-fold serial dilutions in PBS (3 steps). Starting with the OD_600 nm_ 0.2 dilutions, the diluted cultures were applied onto YPAD plates as single 8 μL drops. The plates were incubated at 25-, 30-, and 37 °C and imaged after 3 days.

### UV crosslinking and analysis of cDNA (CRAC)

CRAC was performed using Pol III C128 WT and mutants yeast strains that carried an HTP-tag on subunit C160. The TAP-tag of C128 was, thus, removed for all three tested mutant yeast strains and replaced with a HTP-tag on C160 via homologous recombination. The CRAC protocol used in this study is derived from (Granneman et al., 2009) and modified as previously described (Candelli et al., 2018). Briefly, 2 L of yeast cells expressing an HTP-tagged version of C160 were grown at 30°C to OD_600 nm_ 0.4-0.6 in CSM-TRP medium. Cells were crosslinked for 50 seconds using a W5 UV crosslinking unit (UVO3 Ltd) and harvested by centrifugation. Cell pellets were washed once with cold 1xPBS and resuspended in 2.4 mL/(g of cells) TN150 buffer (50 mM Tris pH 7.8, 150 mM NaCl, 0.1% NP-40 and 5 mM β-mercaptoethanol) in the presence of protease inhibitors (Complete™ EDTA-free Protease Inhibitor Cocktail, Roche). Suspensions were flash frozen in droplets and cells subjected to cryogenic grinding using a Ball Mill MM 400 (5 cycles of 3 minutes at 20 Hz). The resulting frozen cell lysates were thawed on ice, digested with DNase I (165 units per gram of cells) at 25°C for 1h, and then clarified by centrifugation at 13.3 krpm for 30 min at 4°C. Protein extracts were quantified by Bradford reagent (Sigmal) and the same amount of extracts for each sample were subjected to IgG affinity purification as described below.

RNA-protein complexes were bound on M-280 tosylactivated dynabeads coupled with rabbit IgGs (10 mg of beads per sample), washed with TN1000 buffer (50 mM Tris pH 7.8, 1 M NaCl, 0.1% NP-40 and 5 mM β-mercaptoethanol), and eluted by TEV protease digestion. RNAs in the eluates were shortened by treating with 0.2 U of RNase cocktail (RNace-IT, Agilent) and the reaction was stopped by adding of guanidine–HCl to a final concentration of 6 M. The eluates were then incubated with Ni-NTA sepharose (Qiagen, 100 μl of slurry per sample) at 4°C overnight and extensively washed. Sequencing adaptors were ligated to the RNA molecules as described in Granneman et al. (2009). RNA-protein complexes were recovered with elution buffer (50 mM Tris pH 7.8, 50 mM NaCl, 150 mM imidazole, 0.1% NP-40 and 5 mM β-mercaptoethanol) and fractionated by Gel Elution Liquid Fraction Entrapment Electrophoresis (GelFree) system (Expedeon) according to manufacturer’s instructions. The fractions containing Rpc160 were digested with 100 μg of proteinase K, and RNAs were purified and reverse-transcribed using reverse transcriptase Superscript IV (Invitrogen).

The resulting cDNAs were amplified by PCR using LA Taq polymerase (Takara), and the PCR reactions were then treated with 200 U/mL of Exonuclease I (NEB) for 1 h at 37°C. The final library was purified using NucleoSpin columns (Macherey-Nagel) and sequenced on a NextSeq 500 Illumina sequencer.

### CRAC-dataset processing

The sequencing dataset was processed as described in (Xie et al., 2021) with some modification. Briefly, CRAC reads were demultiplexed using the pyBarcodeFilter script from the pyCRACutility suite (Webb et al., 2014). The 5′ adaptor was clipped with Cutadapt and the resulting insert quality-trimmed from the 3′ end using Trimmomatic rolling mean clipping (Bolger et al., 2014). PCR duplicates were collapsed by the pyCRAC script pyFastqDuplicateRemover using a 6-nucleotide random tag included in the 3′ adaptor. The resulting sequences were reverse complemented with the Fastx reverse complement that is included in the fastx toolkit (http://hannonlab.cshl.edu/fastx_toolkit/) and mapped to the R64 genome with bowtie2 using “-N 1” option. Reads shorter than 20 nt were filtered out after mapping. Reads mapped to 37S rDNA unit (transcribed by Pol I) were also excluded from analysis to reduce the background signals. Coverage bigwig files were generated and normalized to counts per million (CPM) using the bamCoverage tool from the deepTools package (Ramírez et al., 2016) using a bin size of 1.

### Bioinformatic analyses

Yeast genome was obtained from Saccharomyces Genome Database (S. cerevisiae genome version R64-2-1). tRNA coordinate files (from 5’ end to the end of the primary terminator) were obtained from (Xie et al., 2021). For each tRNA gene, the primary terminator was defined as the 1st T-tract after the 3’ end of the mature tRNA. Four pairs of tRNAs that are arranged in tandom were excluded from the following analysis.

Reads mapped to different classes of RNAs were summarized by BEDTools coverage. Coverage files were analyzed with deepTools suite (Ramírez et al., 2016). Specifically, for metagene analysis of Pol III occupancy, strand-specific bigwig files and tRNA coordinate files were used as inputs for the computeMatrix tool using a bin size of 1 and the scale-regions mode. The matrices generated were then combined by the computeMatrixOperations tool with the rbind option and used as inputs for the plotProfile tool. For heatmap analyses, the log2 ratio of the Pol III signal in the indicated mutants relative to the WT was calculated by the bigwigCompare tool using a bin size of 1, and the resulting files were used as inputs for the computeMatrix as described above. The final matrix after combination was used as inputs for the plotHeatmap tool. To analyze the correlation between two replicates, the average Pol III signal over regions comprising tRNA genes and 500 bp upstream and downstream regions was computed using the multiBigwigSummary tool. The signals calculated were then compared by plotCorrelation tool to generate scatter plots and compute the correlation coefficients using the Spearman method.

The efficiency of transcription termination in the WT and the mutants was estimated by calculating the read-through index (RT index) defined as the percentage of Pol III signal over the read-through regions (500 bp region immediately downstream of the primary terminator of each tRNA gene) relative to the signal over tRNA gene regions (from 5’ end to the end of the primary terminator). The total Pol III signal at each region was computed with the UCSC bigWigAverageOverBed package (http://genome.ucsc.edu).

Data representation and statistical analyses were performed with R using the ggpubr (https://rpkgs.datanovia.com/ggpubr/) and dplyr (https://dplyr.tidyverse.org/) packages.

---

**Figure S1.**
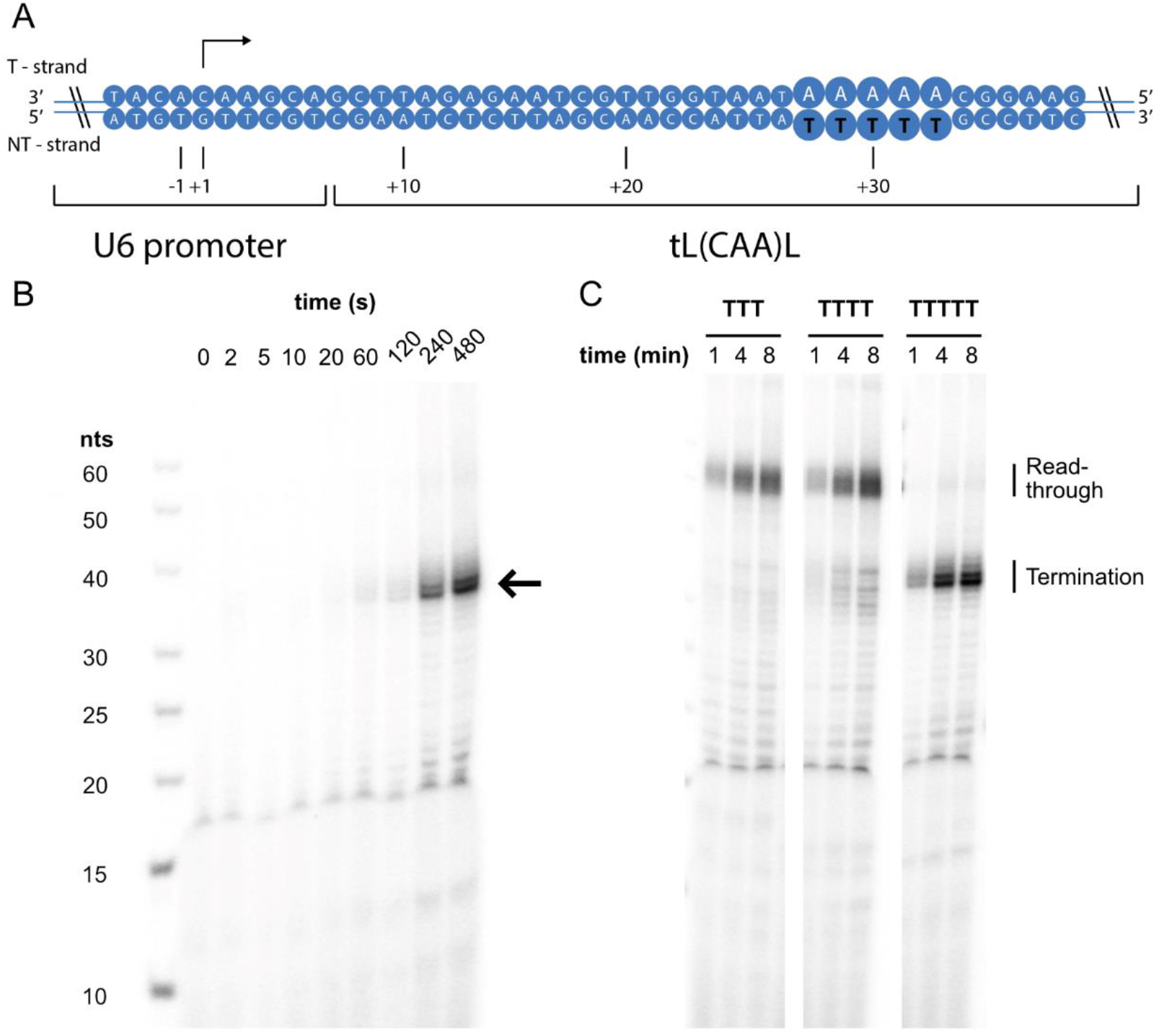
Pol III terminates transcription on poly-dT sequences in vitro. (A) Schematic of the *S. cerevisiae* dsDNA construct, containing the U6 snRNA promoter and the 3’-flanking site of the L(CAA)LR2 gene. The construct was used for promoter-dependent transcription experiments. The termination signal is highlighted in bold letters. (B) Denaturing RNA gel of the promoter-dependent transcription experiment. Shown is a time course of probes taken at the indicated time points. A radioactive RNA product becomes visible over time that is labelled with an arrow; n (dsDNA) = 2.5 pmol; n (Pol III) = 5 pmol; n (TFIIIB) = 5 pmol. 15% TBE denaturing PAGE. (C) Time-course of the Pol III transcription experiment testing the effect of a 5-dT, 4-dT, or 3-dT sequence on Pol III transcription termination. Same conditions as in (B). The gel has been cropped for clarity.

**Figure S2.**
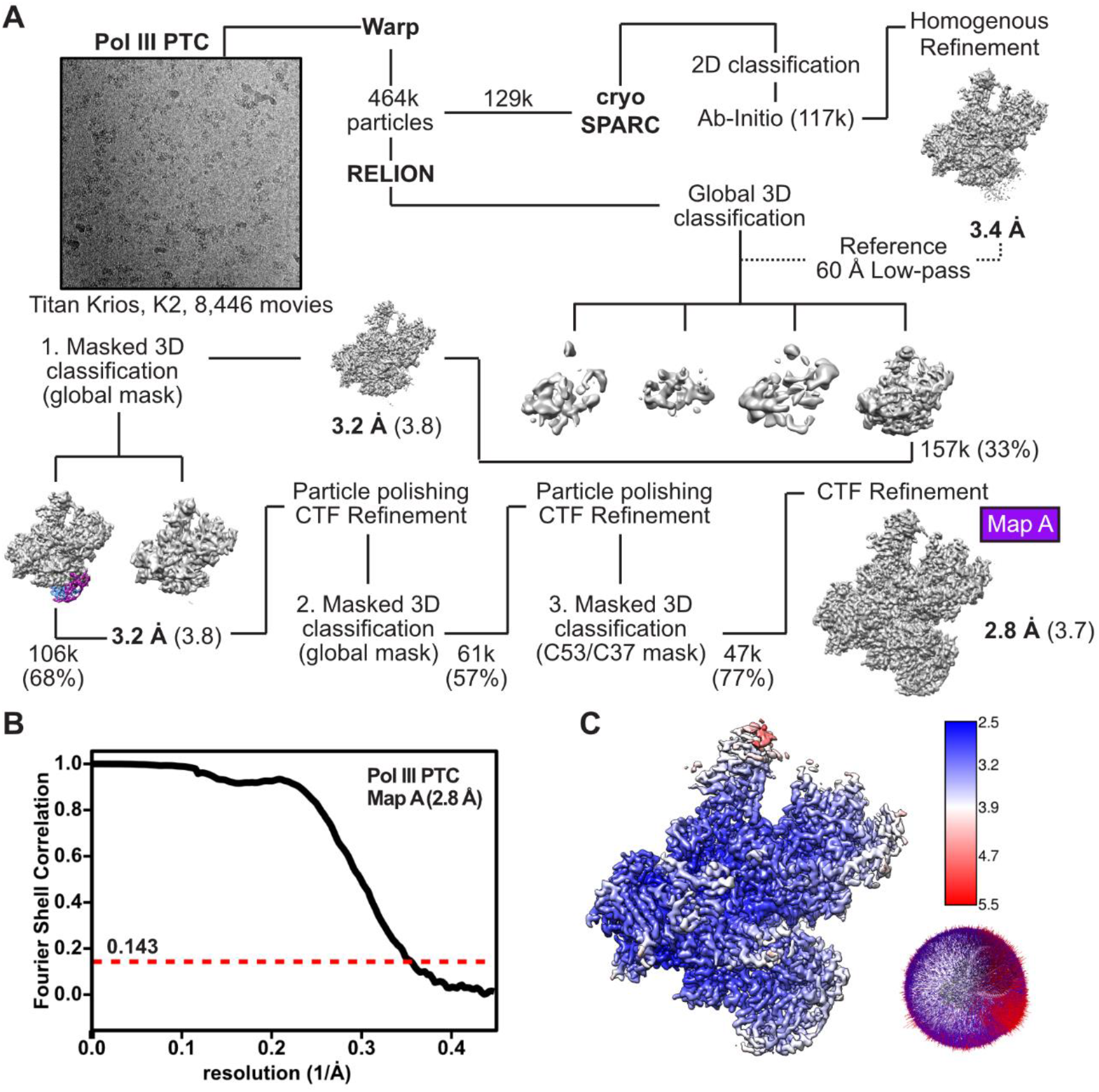
Data-collection and -processing workflow of the yeast Pol III PTC. (A) Cryo-EM data collection and processing strategy for the yeast Pol III PTC. Particle numbers are rounded down. Shown are the unsharpened cryo-EM maps obtain via 3D refinement in RELION. Reported resolution values are given below the cryo-EM maps and correspond to sharpened maps obtained via RELION post-processing. The resolution values of the unsharpened 3D maps are added in round brackets. Map features that show improved cryo-EM signal after masked 3D classification are highlighted in colours. (B) Fourier shell correlation (FSC) curve of the yeast Pol III PTC derived via RELION post-processing (FSC = 0.143). (C) Estimation of local resolution and angular distribution plot of the yeast Pol III PTC. Shown is the unsharpened cryo-EM map.

**Figure S3.**
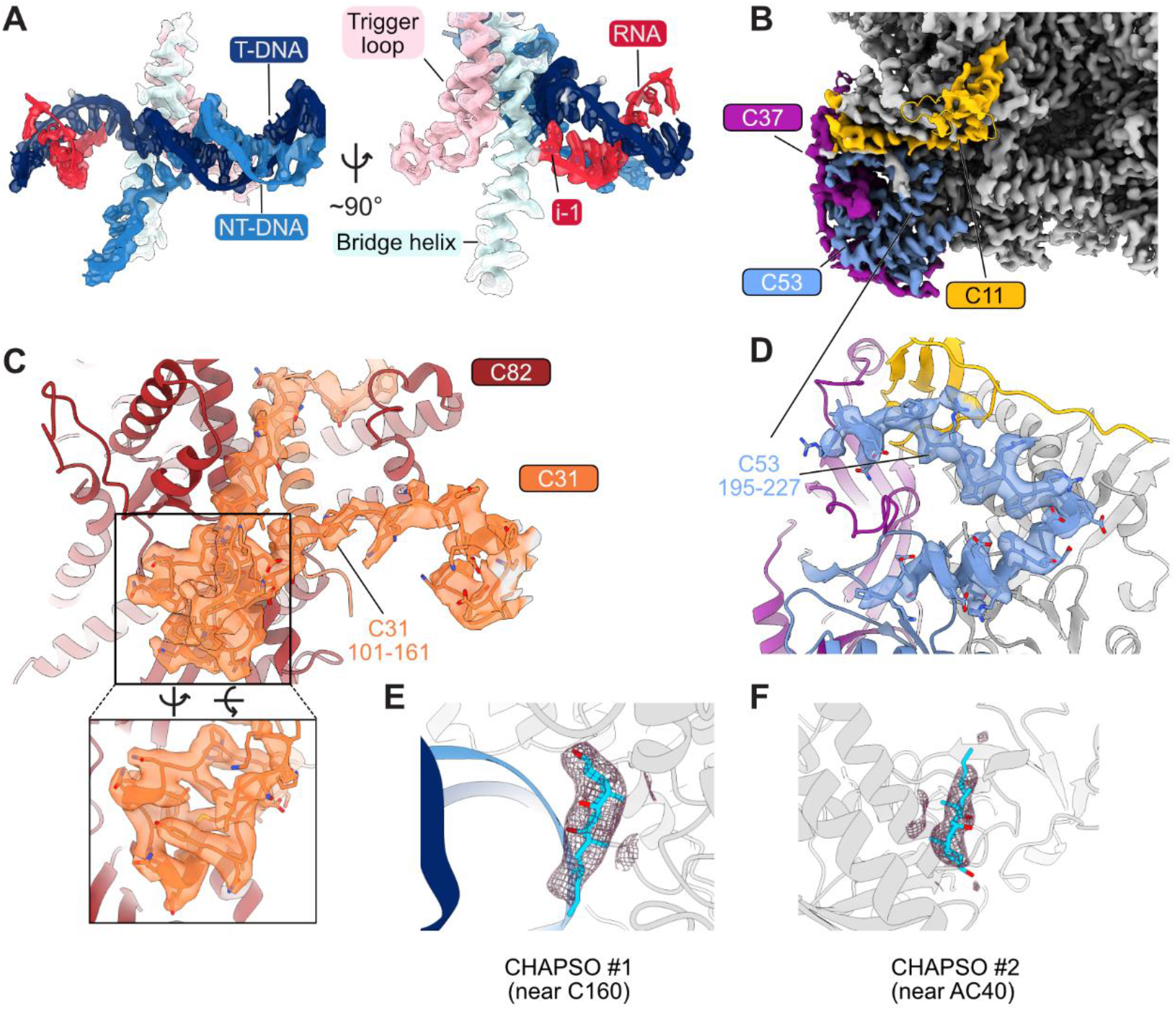
Cryo-EM data quality of the active site elements and newly build regions. (A) Close-up views on the Pol III PTC active site elements. Cryo-EM densities are shown as transparent surfaces. (B) The C11 C-terminal domain is visible and binds the Pol III core close to the C160 jaw domain and subunit ABC27. (C) Newly built region (101-161) of the Pol III heterotrimer subunit C31. Cryo-EM density of the newly assigned region is shown as a transparent surface. (D) Newly built region (195-227) of the Pol III heterodimer subunit C53. Cryo-EM density of the newly assigned region is shown as a transparent surface. (E) - (F) Cryo-EM densities of putative CHAPSO molecules, of which the tetracyclic sterol head groups were placed.

**Figure S4.**
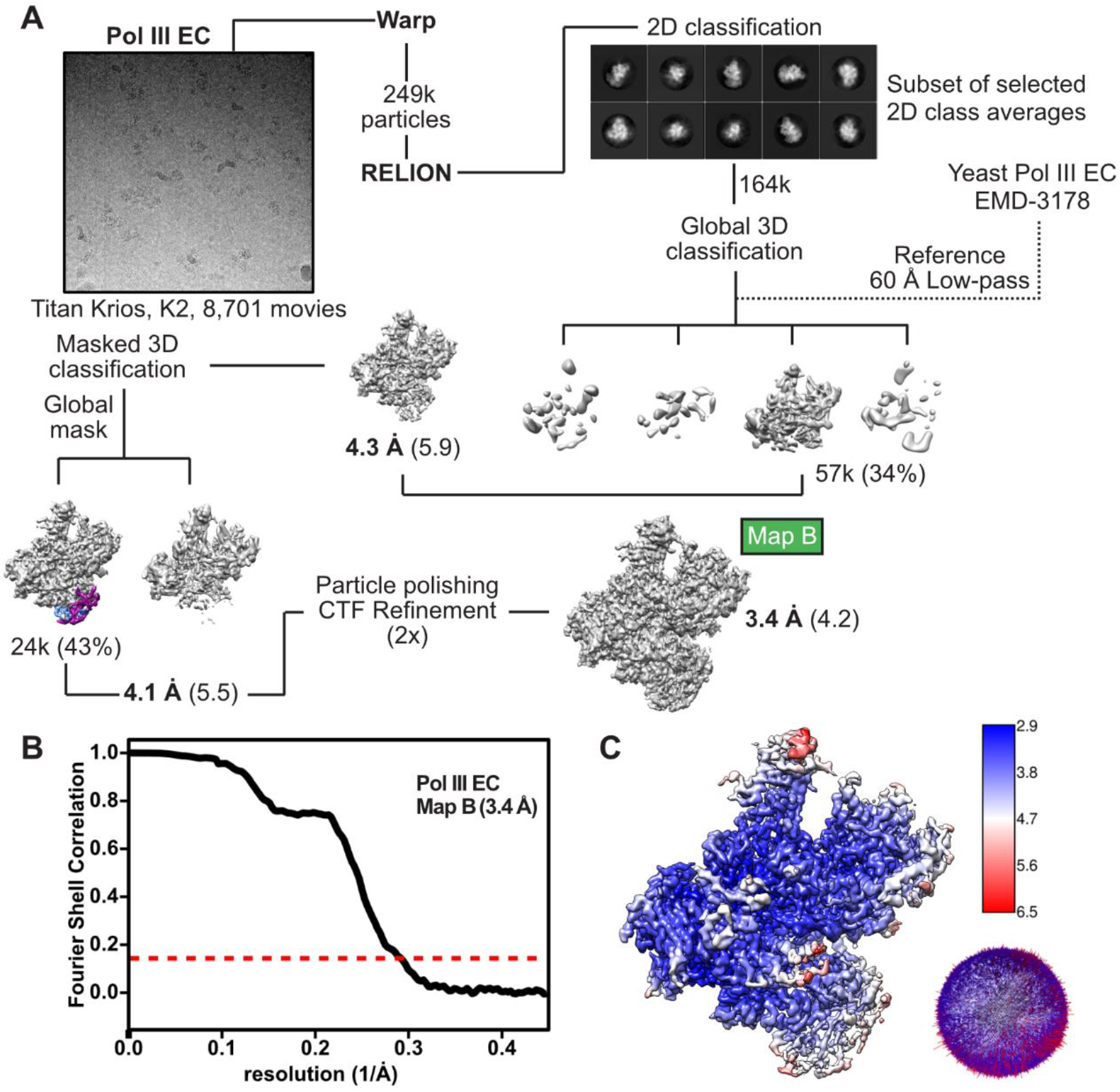
Data-collection and -processing workflow of the yeast Pol III EC. (A) Cryo-EM data collection and processing strategy for the yeast Pol III EC. Particle numbers are rounded down. Shown are the unsharpened cryo-EM maps obtain via 3D refinement in RELION. Reported resolution values are given below the cryo-EM maps and correspond to sharpened maps obtained via RELION post-processing. The resolution values of the unsharpened 3D maps are added in round brackets. Map features that show improved cryo-EM signal after masked 3D classification are highlighted in colours. (B) Fourier shell correlation (FSC) curve of the yeast Pol III EC derived via RELION post-processing (FSC = 0.143). (C) Estimation of local resolution and angular distribution plot of the yeast Pol III EC. Shown is the unsharpened cryo-EM map.

**Figure S5.**
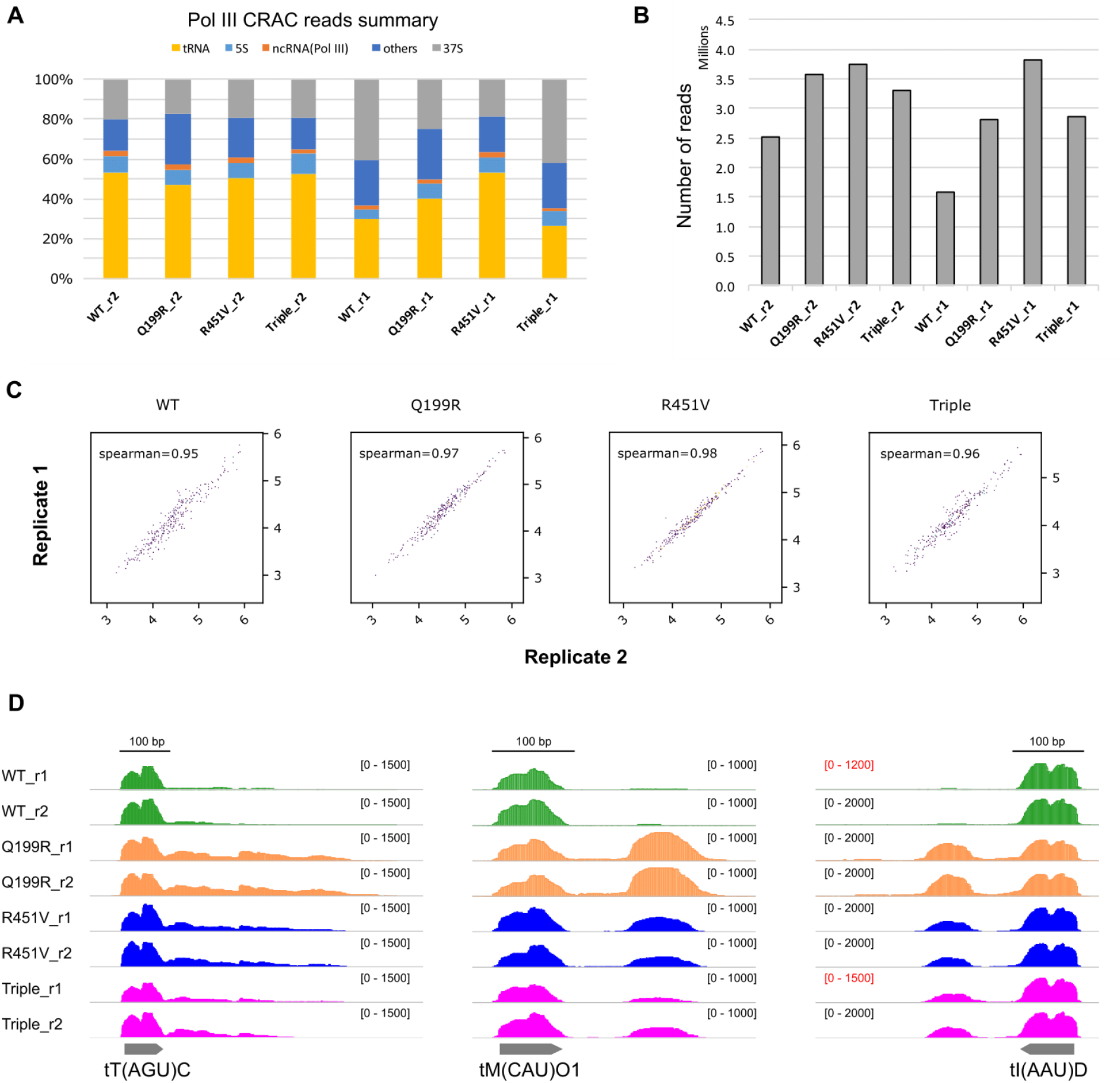
Complementary analyses validating CRAC experiments in Figures 4. A) Comparison of the read distribution among different genomic regions in the indicated strains in our Pol III CRAC experiment. The “others” category corresponds to Pol II genes and intergenic regions. Note that we have more contaminants and lower signal at tRNA regions in the WT_r1 and Triple_r1 samples, which, however, does not significantly affect the assessment of the transcription termination efficiency. B) Plot representing the number of mapped reads obtained in a typical CRAC experiment in the different samples. “M” denotes millions. C) Scatter plots showing the high correlation, calculated by the Spearman method, between the two biological replicates of each strain for the CRAC experiments showed in figure 4. D) Integrative Genomics Viewer (IGV) screenshots of examples of tRNA genes displaying termination defects in the two biological replicates of each strains. The values under brackets correspond to the scale of the Pol III signal expressed in CPM. Note that because of the higher amounts of contaminants in the WT_r1 and Triple_r1 samples, the signal in the tRNA regions is lower (scale indicated in red), but the Pol III distribution is similar as in the other replicate of the same strains (r2).

**Figure S6.**
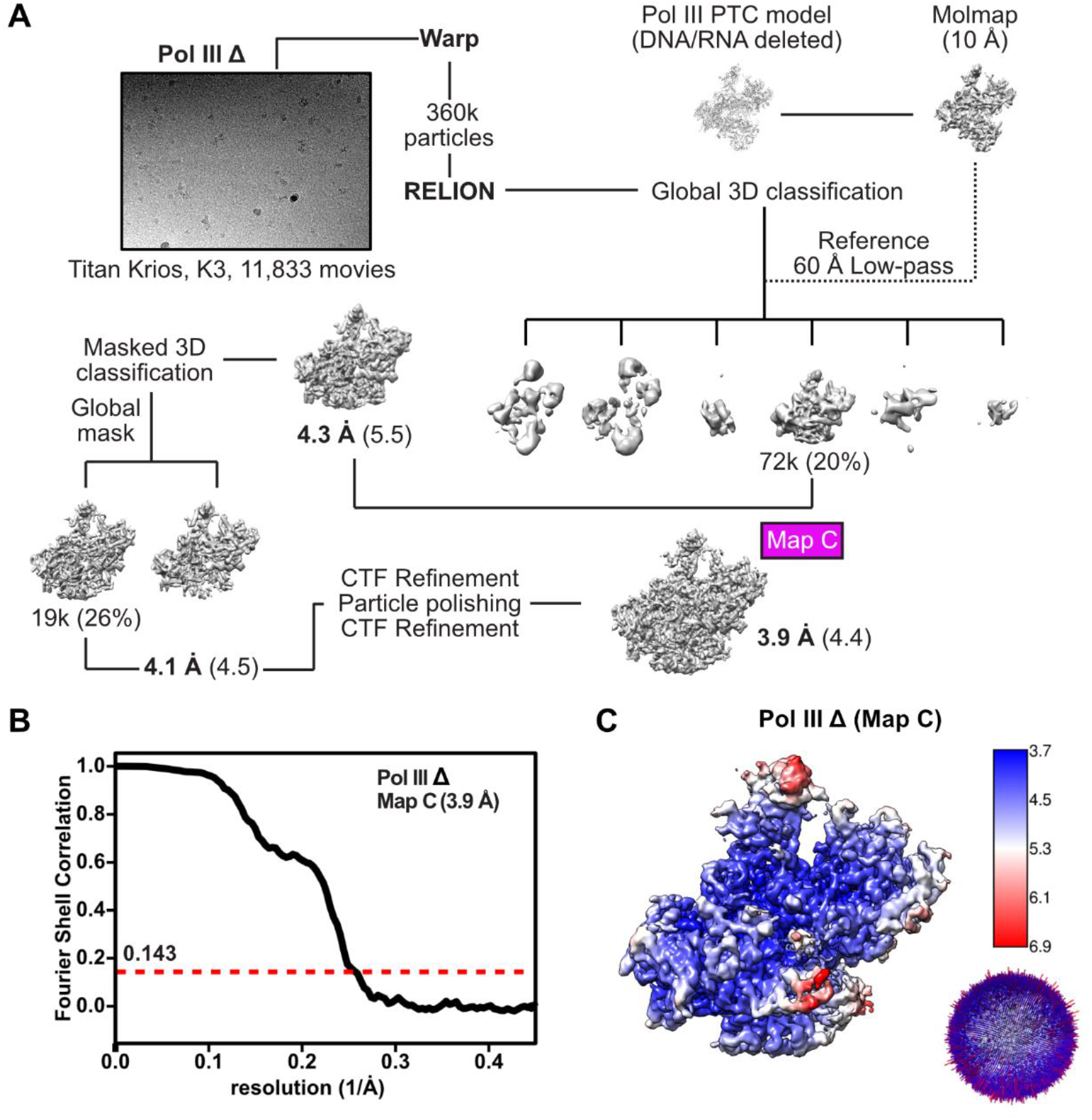
Data-collection and -processing workflow of yeast Pol III Δ. (A) Cryo-EM data collection and processing strategy for the yeast Pol III Δ. Particle numbers are rounded down. Shown are the unsharpened cryo-EM maps obtain via 3D refinement in RELION. Reported resolution values are given below the cryo-EM maps and correspond to sharpened maps obtained via RELION post-processing. The resolution values of the unsharpened 3D maps are added in round brackets. (B) Fourier shell correlation (FSC) curve of the yeast Pol III Δ derived via RELION post-processing (FSC = 0.143). (C) Estimation of local resolution and angular distribution plot of the yeast Pol III Δ. Shown is the unsharpened cryo-EM map.

**Figure S7.**
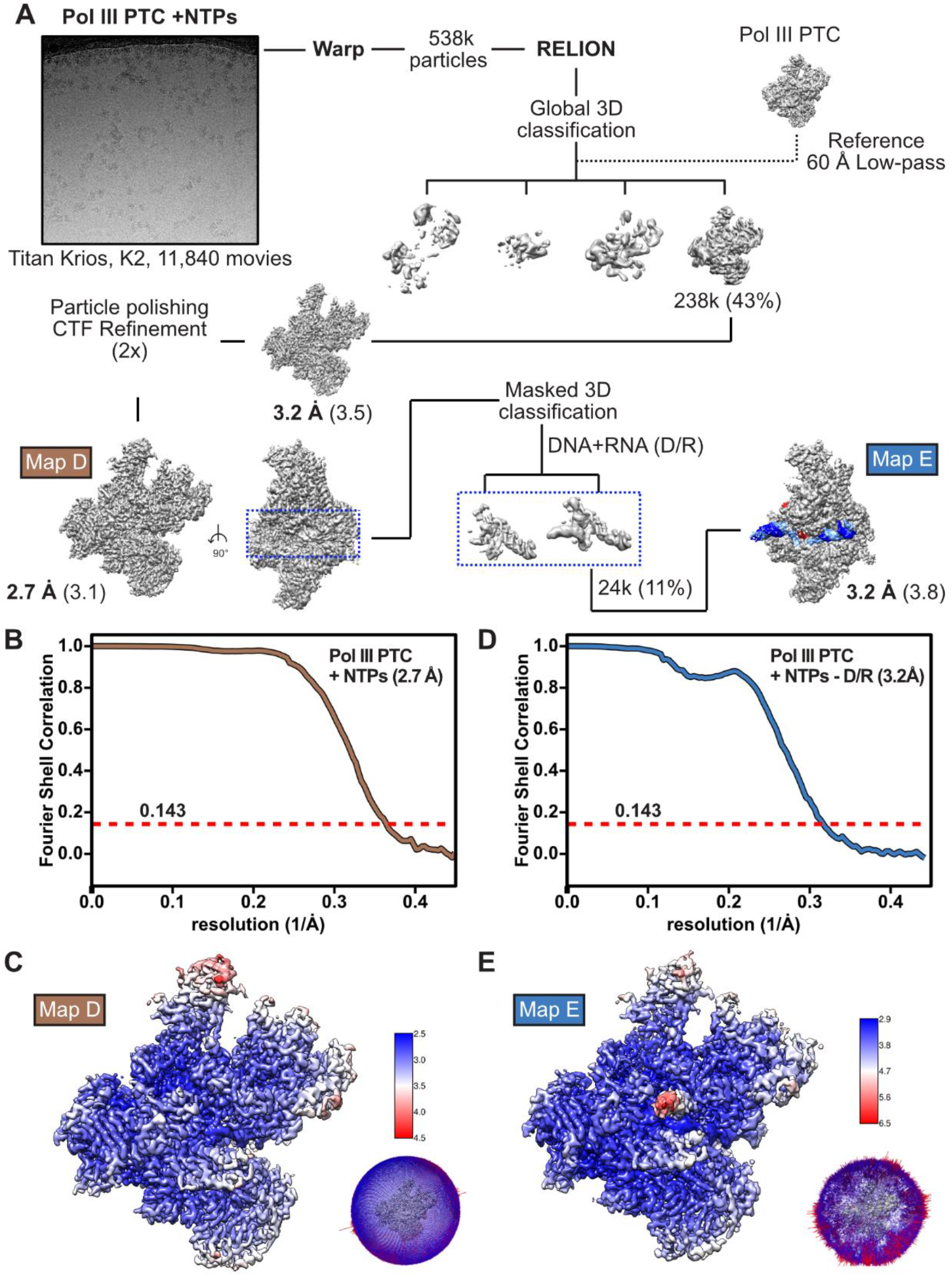
Data-collection and -processing workflow of yeast Pol III PTC + NTPs. (A) Cryo-EM data collection and processing strategy for the yeast Pol III PTC + NTPs. Particle numbers are rounded down. Shown are the unsharpened cryo-EM maps obtain via 3D refinement in RELION. Reported resolution values are given below the cryo-EM maps and correspond to sharpened maps obtained via RELION post-processing. The resolution values of the unsharpened 3D maps are added in round brackets. Map features that show improved cryo-EM signal after masked 3D classification are highlighted in colours. (B) Fourier shell correlation (FSC) curve of the yeast Pol III PTC + NTPs derived via RELION post-processing (FSC = 0.143). (C) Fourier shell correlation (FSC) curve of the yeast Pol III PTC + NTPs map that was subjected to masked 3D-classification on the bound DNA/RNA (D/R) elements. (D) Estimation of local resolution and angular distribution plot of the yeast Pol III PTC + NTPs. Shown is the unsharpened cryo-EM map. (E) Estimation of local resolution and angular distribution plot of the yeast Pol III PTC + NTPs classified on the DNA/RNA elements. Shown is the unsharpened cryo-EM map.

**Table S1.** Cryo-EM data collection, refinement and validation statistics.

|  | **Pol III**<br>**PTC**<br>**Map A** | **Pol III**<br>**EC**<br>**Map B** | **Pol III**<br>**$\Delta$**<br>**Map C** | **Pol III PTC**<br>**+ NTPs**<br>**Map D** | **Pol III PTC**<br>**+ NTPs 2**<br>**Map E** |
| --- | --- | --- | --- | --- | --- |
| PDB code | 7Z1L | 7Z1M | 7Z1N | 7Z1O | - |
| EMDB code | EMD-14447 | EMD-14448 | EMD-14449 | EMD-14451 | EMD-14450 |
| **Data collection and processing** |  |  |  |  |  |
| Magnification | 130,000 | 130,000 | 81,000 | 130,000 | 130,000 |
| Voltage (kV) | 300 | 300 | 300 | 300 | 300 |
| Electron exposure (e-/Å<sup>2</sup>) | 40.9 | 38.1 | 39.6 | 44.8 | 44.8 |
| Defocus range (μm) | 0.75-2.25 | 0.50-1.75 | 1.00-2.50 | 0.75-2.25 | 0.75-2.25 |
| Pixel size (Å) | 1.041 | 1.040 | 1.053 | 1.041 | 1.041 |
| Symmetry imposed | C1 | C1 | C1 | C1 | C1 |
| Initial particle images (no.) | 464,321 | 248,884 | 359,678 | 538,161 | 538,161 |
| Final particle images (no.) | 47,222 | 24,295 | 18,530 | 237,468 | 23,641 |
| Map resolution (Å) | 2.8 | 3.4 | 3.9 | 2.7 | 3.2 |
| FSC threshold | 0.143 | 0.143 | 0.143 | 0.143 | 0.143 |
| Map resolution range (Å) | 2.5-8.0 | 3.2-8.8 | 3.7-8.5 | 2.5-5.1 | 2.9-7.5 |
| **Refinement** |  |  |  |  |  |
| Initial model used (PDB code) | 6TUT, 5FJ8,<br>5FJA | 6TUT, 5FJ8,<br>5FJA | 6TUT, 5FJ8,<br>5FJA | 6TUT, 5FJ8,<br>5FJA |  |
| Model resolution (Å) | 3.0 | 3.4 | 3.9 | 3.0 |  |
| FSC threshold | 0.5 | 0.5 | 0.5 | 0.5 |  |
| Map sharpening *B* factor (Å<sup>2</sup>) | -27.6 | -70.4 | -107.1 | -66.4 |  |
| **Model composition** |  |  |  |  |  |
| Non-hydrogen atoms | 42,330 | 42,127 | 38,605 | 42,375 |  |
| Protein/nucleotide residues | 5156/63 | 5153/53 | 4705/56 | 5154/66 |  |
| Ligands | 5x Zn, 1x Mg,<br>2x 1N7 | 5x Zn, 1x Mg,<br>2x 1N7 | 5x Zn, 1x Mg,<br>2x 1N7 | 5x Zn, 1x Mg,<br>2x 1N7 |  |
| ***B* factors (Å<sup>2</sup>)** |  |  |  |  |  |
| Protein | 73.63 | 112.52 | 56.95 | 61.03 |  |
| Nucleotide | 90.42 | 157.39 | 84.51 | 79.65 |  |
| Ligand | 73.44 | 116.06 | 61.34 | 67.07 |  |
| **R.m.s. deviations** |  |  |  |  |  |
| Bond lengths (Å) | 0.003 | 0.004 | 0.005 | 0.003 |  |
| Bond angles (°) | 0.636 | 0.678 | 0.779 | 0.640 |  |
| **Validation** |  |  |  |  |  |
| MolProbity score | 1.34 | 1.45 | 1.68 | 1.28 |  |
| Clashscore | 4.14 | 4.73 | 6.26 | 3.96 |  |
| Poor rotamers (%) | 0.02 | 0.00 | 0.00 | 0.00 |  |
| **Ramachandran plot** |  |  |  |  |  |
| Favored (%) | 97.23 | 96.68 | 95.13 | 97.55 |  |
| Allowed (%) | 2.77 | 3.30 | 4.83 | 2.45 |  |
| Disallowed (%) | 0.00 | 0.02 | 0.04 | 0.00 |  |

